# CryoEM reveals unprecedented binding site for Na_V_1.7 inhibitors enabling rational design of potent hybrid inhibitors

**DOI:** 10.1101/2022.11.10.515983

**Authors:** Marc Kschonsak, Christine C. Jao, Christopher P. Arthur, Alexis L. Rohou, Philippe Bergeron, Daniel Ortwine, Steven J. McKerrall, David H. Hackos, Lunbin Deng, Jun Chen, Peter S. Dragovich, Matthew Volgraf, Matthew R. Wright, Jian Payandeh, Claudio Ciferri, John C. Tellis

**Author notes:** Corresponding authors. (J.P); (C.C.); (J.C.T.). **Author Contributions:** J.P., C.C., and J.C.T. designed research, M.K., C.J., C.P.A., A.R., D.H., and J.C., performed research, J.C.T, P.B., D.O., S.M., P.S.D., and M.V. analyzed data and contributed to compound design. J.C.T., C.C., J.P., and M.K. wrote the paper. **Competing Interest Statement:** The authors declare no competing financial interest.

## Abstract

The voltage-gated sodium (NaV) channel NaV1.7 has been identified as a potential novel pain target due to its striking human genetics. However, clinically available drugs (e.g. lidocaine, carbamazepine, etc.) are not selective among the nine NaV channel subtypes, NaV1.1-NaV1.9, and the two currently known classes of NaV1.7 subtype-selective inhibitors (aryl- and acylsulfonamides) have undesirable characteristics that may limit their development. Moreover, understanding of the structure-activity relationships of the acylsulfonamide class of NaV1.7 inhibitors, exemplified by the clinical development candidate **GDC-0310**, has been based solely on a single co-crystal structure of an arylsulfonamide inhibitor series. To advance inhibitor design targeting the NaV1.7 channel, we established an iterative system to routinely obtain high-resolution ligand-bound NaV1.7 structures using cryogenic electron microscopy (cryo-EM). We report that **GDC-0310** engages the NaV1.7 voltage-sensing domain 4 (VSD4) through an unexpected binding mode orthogonal to the arylsulfonamide class binding pose, which identifies a previously unknown ligand binding site in NaV channels. This finding enabled the design of a novel hybrid inhibitor series that bridges the aryl and acylsulfonamide binding pockets and allows for the generation of molecules with substantially differentiated structures and properties. Overall, this study highlights the power of cryo-EM methods to pursue challenging drug targets using iterative and high-resolution structure-guided inhibitor design. It also underscores an important role of the membrane bilayer in the discovery of selective NaV channel modulators.

## Introduction

Voltage-gated sodium (NaV) channels initiate and propagate action potentials in excitable tissues and play important roles in health and disease.^1,2^ The NaV1.7 channel is expressed predominantly in the peripheral nervous system and genetic studies have identified compelling loss-of-function and gain-of-function phenotypes in human pain syndromes, prompting significant efforts to develop NaV1.7-selective inhibitors as potential novel analgesic drugs.^3-5^ NaV channels contain 24-transmembrane segments linked in four homologous domains (DI-DIV), where four peripheral voltage-sensor domains (VSD1-4) surround a central ion-conducting pore module that houses the ion selectivity filter and key ligand and toxin receptor sites. Traditionally, all clinically available NaV channel inhibitors lack significant molecular selectivity among the NaV1.1– 1.9 subtypes owing to the high sequence conservation found at the ligand binding site within the central cavity of the ion conducting pore module.^5,6^

A breakthrough study in 2013 by McCormack and colleagues reported the discovery of an arylsulfonamide antagonist, **PF-04856264**, that bound to an unprecedented receptor site in VSD4 with demonstrated molecular selectivity for human NaV1.7 over other subtypes.^7^ However, the related development candidate **PF-05089771** did not meet clinical endpoints in human subjects with painful diabetic peripheral neuropathy, possibly due to poor target coverage and the intolerable doses required to achieve efficacy.^8-10^ Additionally, an alternate acylsulfonamide inhibitor that also targeted the VSD4 receptor site in NaV1.7, **GDC-0276**, was halted in Phase I clinical trials due to safety concerns and potential off-target effects likely attributed to the high lipophilicity of the compound.^11,12^ To date, selective NaV1.7 inhibitors with an improved therapeutic index relative to these clinical-stage compounds have not yet been identified, and this absence underscores the need to optimize such molecules using modern, structure-guided design approaches.

The inherent complexity and dynamic nature of human NaV channels have historically presented significant barriers to obtaining high-resolution experimental structural information, especially of inhibitor-bound complexes.^13^ Using an engineered human VSD4 NaV1.7-NaVAb bacterial channel chimera and X-ray crystallography, the binding mode of the **GX-936** arylsulfonamide inhibitor (Supplementary Figure 1) revealed that the anionic sulfonamide group engages the fourth arginine gating charge (R4) to trap VSD4 in an activated conformation, which stabilizes a non-conductive, inactivated state of the channel.^14^ While determinants of subtype selectivity and structure-activity relationships (SAR) of the arylsulfonamide inhibitor class could be rationalized by the **GX-936** co-crystal structure, additional ligand-bound structures of suitable resolution were not returned by the NaV1.7-NaVAb chimeric channel crystallography system. This shortcoming led molecular docking studies to presume that the acylsulfonamide inhibitors also bound VSD4 in an analogous manner, despite points of inexplicable SAR.^15^ The overall paucity of direct structural information has made the optimization of NaV1.7 inhibitors challenging, with many questions about the determinants of potency, selectivity, and the relationship between the acylsulfonamide and arylsulfonamide inhibitor classes remaining unanswered.

Over the last three decades, protein crystallography and structure-based drug design (SBDD) have become gold standards across the pharmaceutical industry for the identification of ligand binding pockets and the optimization of drug candidates for clinical development. While SBDD has proven successful for many important targets, including G protein-coupled receptors (GPCRs),^16^ its applicability to several membrane protein targets, such as NaV channels, has been limited due to the extreme difficulties with their iterative crystallization and structure determination.^13^ Cryogenic electron microscopy (cryo-EM) has recently emerged as a transformative technique to determine the high-resolution structures of diverse protein targets and has proven particularly effective for membrane proteins that are recalcitrant to crystallization.^17^ Despite this breakthrough, cryo-EM structure determination of membrane targets in complex with small or large molecule therapeutics frequently remains retrospective and is often enabled after the advancement of key molecules into the clinic.^18,19^ Sluggish turn-around times and modest resolutions typically afforded by cryo-EM have limited its application for real-time SBDD efforts. Here, we describe a system for the iterative determination of high-resolution NaV1.7 small molecule co-structures via cryo-EM that has led to the development of a novel class of inhibitors. Critical to this advancement has been establishment of a robust protocol for sample preparation and structure determination and the enablement of the first structure of an acylsulfonamide bound to NaV1.7 VSD4, revealing a previously unknown pocket between the S3 and S4 helices. Consequently, a novel hybrid inhibitor series bridging the aryl and acylsulfonamide pockets was generated. Our work exemplifies the deployment of cryo-EM as a workhorse structural biology tool in an active medicinal chemistry campaign and thus represents an important milestone toward NaV channel drug discovery.

## Results

### Structure of the G-3565-VSD4 NaV1.7-NaVPas channel complex in lipid nanodiscs

Our approach to establish an iterative, high-resolution system to enable NaV1.7 SBDD was ultimately guided by three observations: our repeated failure to generate well-diffracting crystals of the NaV1.7-NaVAb bacterial channel chimera system,^14^ our difficulty of reproducibly expressing a suitable amount of full-length human NaV1.7 channel,^20,21^ and the very limited local resolution observed at VSD4 in all available human Nav1.7 cryo-EM structures.^22,23^ Thus, we sought to exploit an engineered human VSD4-NaV1.7-NaVPas cockroach channel chimeric construct that had been previously shown to complex with small molecule inhibitors and peptide toxins known to target VSD4.^21^ The robust protein yields following expression and purification allowed us to readily pursue reconstitution of the VSD4-NaV1.7-NaVPas channel into lipid nanodisc with high recovery (Supplementary Figure 2). Small molecule inhibitors were added prior to sample vitrification, followed by cryo-EM data collection and processing procedures, which allowed us to routinely obtain 3D-reconstructions in the 2.2-3.0 Å resolution range (Figure 1, Supplementary Figure 3).

**Figure 1:**
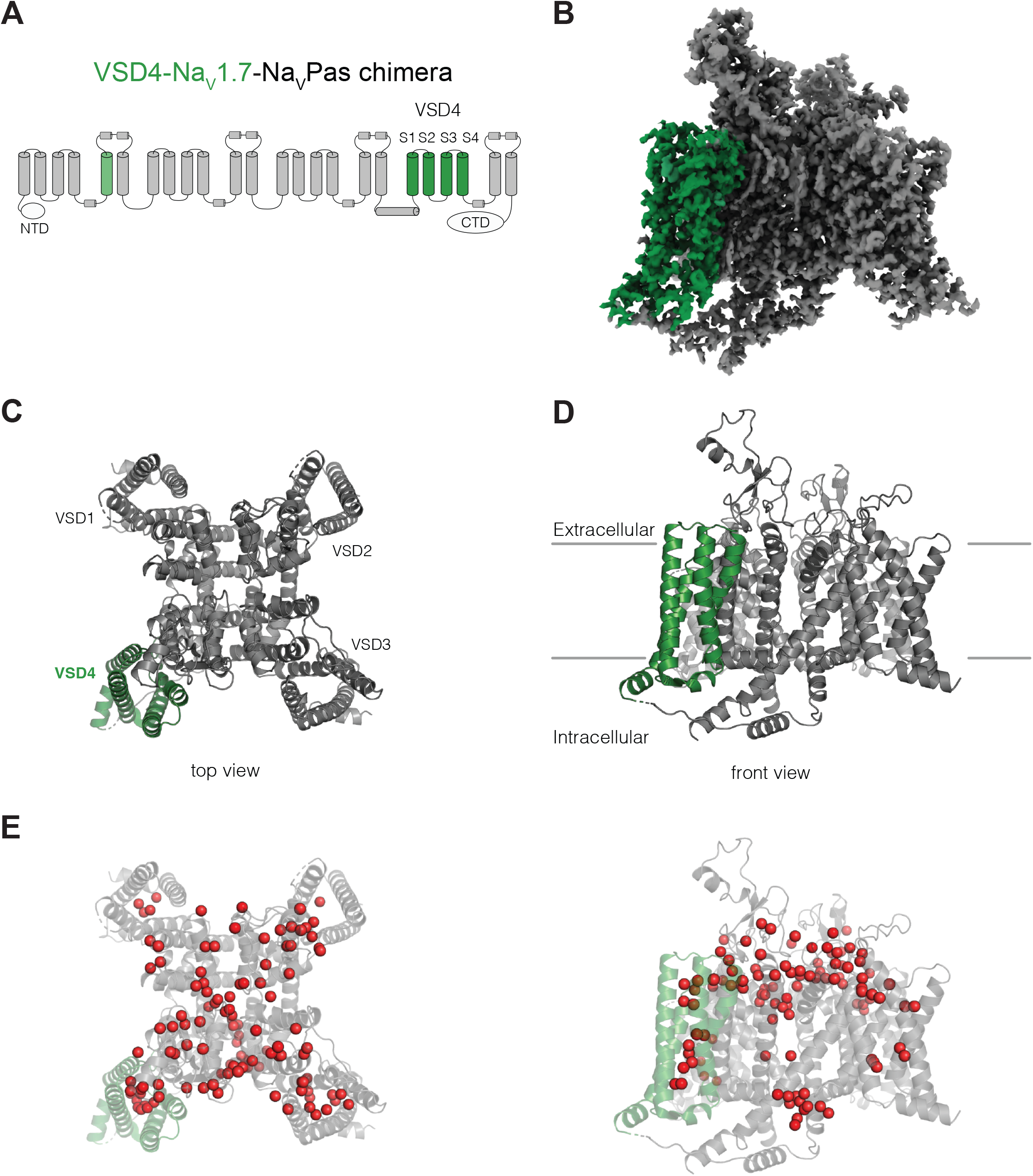
**(A)** Schematic of the VSD4-NaV1.7-NaVPas channel. The portions humanized to the NaV1.7 sequence are shown in green. N-terminal domain (NTD) and CTD are indicated. **(B)** Side view of the single-particle cryo-EM reconstruction of VSD4-NaV1.7-NaVPas channel **(C-D)** Cartoon representations of the Top and Side views of VSD4-NaV1.7-NaVPas channel. Individual VSD domains are indicated. VSD4 is highlighted in green. **(E)** Localization of water molecules (in red) in the VSD4-NaV1.7-NaVPas channel structure. VSD4 is highlighted in green.

The VSD4-NaV1.7-NaVPas chimera displays the expected domain-swapped arrangement with numerous densities assigned as phospholipids bound to the channel, confirming the maintenance of a native membrane-like environment (Figure 1). As seen in previous NaVPas structures,^21,24,25^ the ion-conducting pore module of the VSD4-NaV1.7-NaVPas chimeric channel is closed, consistent with a nonconductive or inactivated state. The quality and resolution of our structure allowed us to assign 112 water molecules for the highest resolution structure (Figure 1, Video 1). The visualization of many well-resolved water molecules bound within the VSDs and coordinated within the ion selectivity filter provides new insights into the interactions that might occur during gating charge transfer and ion conduction, respectively.

Although we were able to determine the crystal structure of VSD4-NaVAb bound to the arylsulfonamide inhibitor **GX-936**,^14^ no well-diffracting crystals were obtained for any other arylsulfonamide compounds despite years of continued effort. Arylsulfonamide **GNE-3565** is representative of an advanced series of arylsulfonamide class NaV1.7 inhibitors that demonstrates channel blockage at subnanomolar concentrations with mixed subtype selectivity (Figure 2A). We complexed **GNE-3565** with the VSD4 NaV1.7-NaVPas channel-nanodisc system and employed cryo-EM to assess its overall binding pose (Figure 2B, C). The 2.9 Å resolution cryo-EM map revealed that the S1-S2 and S3-S4 helices from VSD4 form a clamshell-like structure that closes over **GNE-3565** (Figure 2D, E, Supplementary Data Figure 4A-D, Supplementary Data Table 1), similar to the binding pose reported for **GX-936**.^14^ Specifically, the ionized arylsulfonamide group of **GNE-3565** (measured pKa = 5.8) bisects VSD4 to salt-bridge directly to R4 on the S4 helix, while the central phenyl ring extends perpendicular between the S2 and S3 helices to directly contact established subtype-selectivity determinants Tyr1537 and Trp1538 on the S2 helix (Figure 2A-D). As for **GX-936**, the **GNE-3565** arylsulfonamide VSD4 receptor binding site can be divided into three regions: an anion-binding pocket, a selectivity pocket, and a lipid-exposed pocket (Figure 2D). The root-mean-square deviation (RMSD) between the **GNE-3565**-VSD4 and the **GX-936**-VSD4 structures^14^ is 0.667 (714 to 714 atoms), (Supplementary Figure 5A), supporting long-held assumptions by medicinal chemistry teams that all structurally-related arylsulfonamides should be expected to complex to VSD4 through similar determinants.^8,26-31^

**Figure 2:**
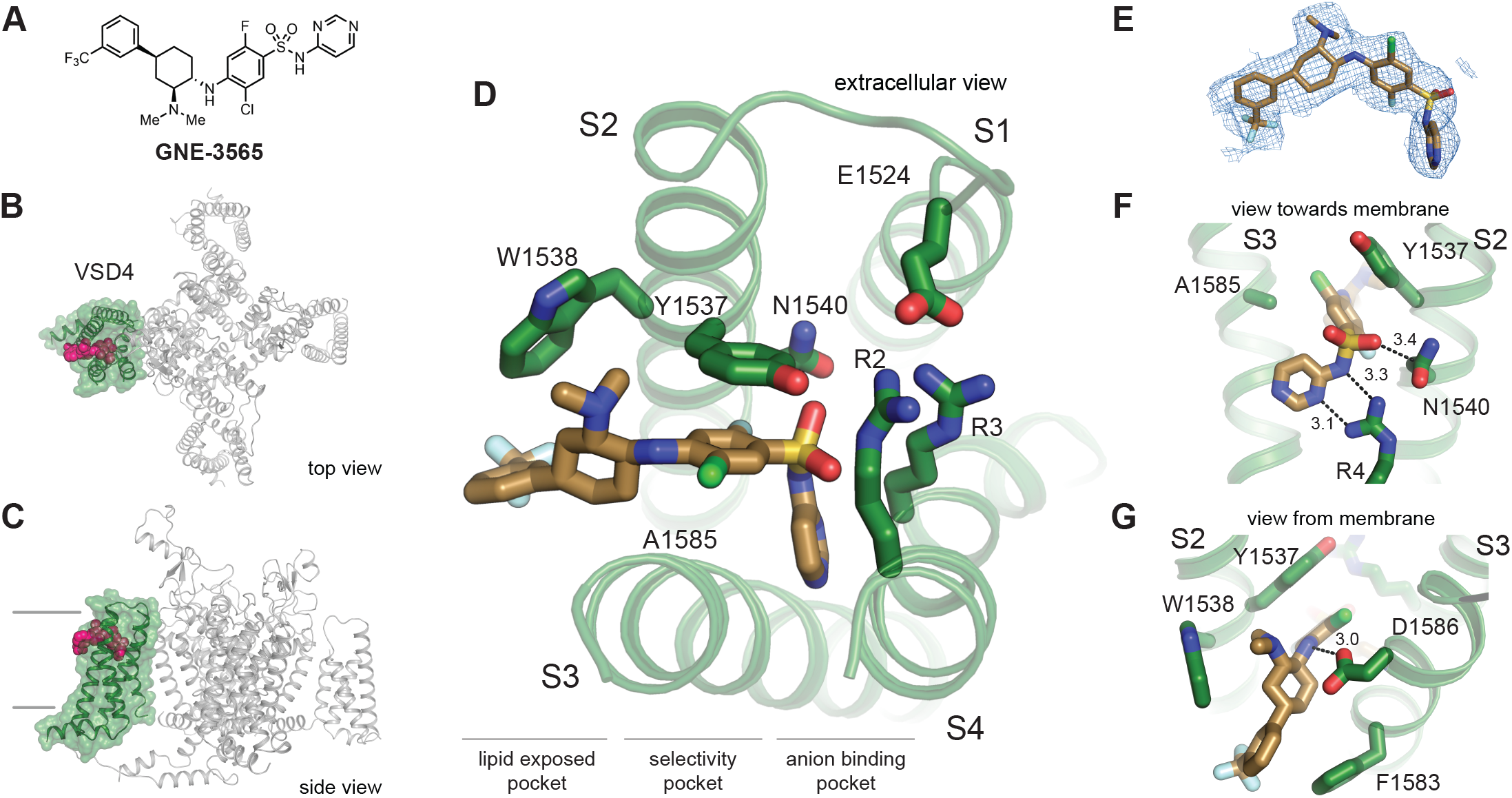
**(A)** Chemical structures of arylsulfonamide **GNE-3565. (B**,**C)** Top and Side views of VSD4-NaV1.7-NaVPas channel bound to **GNE-3565**. VSD4 is highlighted in green, **GNE-3565** in magenta. **(D)** Extracellular view of VSD4-NaV1.7-NaVPas arylsulfonamide receptor site is shown with select side chains rendered as sticks. **(E)** The cryoEM map surrounding the ligand **GNE-3565** is shown in mesh representation. **(F)** View towards the membrane highlighting key interactions with the **GNE-3565** warhead. **(G)** View from the membrane highlighting key interactions with the **GNE-3565** central phenyl ring.

### Structure of the acylsulfonamide GDC-0310 in complex with the VSD4 NaV1.7-NaVPas channel

To investigate the binding pose of a representative NaV1.7 selective acylsulfonamide, **GDC-0310** (Figure 3A) was complexed with the VSD4 NaV1.7-NaVPas channel in nanodiscs and a cryo-EM structure was determined to 2.5 Å resolution (Figure 3B-D, Supplementary Data Figure 3G-J, Supplementary Data Table 1). Remarkably, the binding mode of **GDC-0310** revealed an unexpected and previously unknown pocket that formed between the S3 and S4 helices (Figure 3B-E). While the aryl and acyl moieties of **GDC-0310** and **GNE-3565** bind in the same region, the remainder of the acyl receptor site is orthogonal to the corresponding binding domain reported for the aryl class (Figure 3B-D). In detail, the anionic acylsulfonamide of **GDC-0310** participates in salt-bridge bonding interactions with R4 and R3 through the carbonyl and sulfonamide oxygen atoms, respectively (Figure 3F). Moreover, the aryl ring of the **GDC-0310** splits the S3 and S4 helices to occupy a lipophilic pocket, displacing the S3 helix laterally by ∼3 Å relative to the **GNE-3565**-VSD4 structure (Figure 3G). Our cryo-EM inhibitor bound structures serve to highlight a dynamic environment in VSD4 where the S1-S4 helixes can be differentially separated by distinct small molecule binding poses (Figure 2D, Figure 3D).

**Figure 3:**
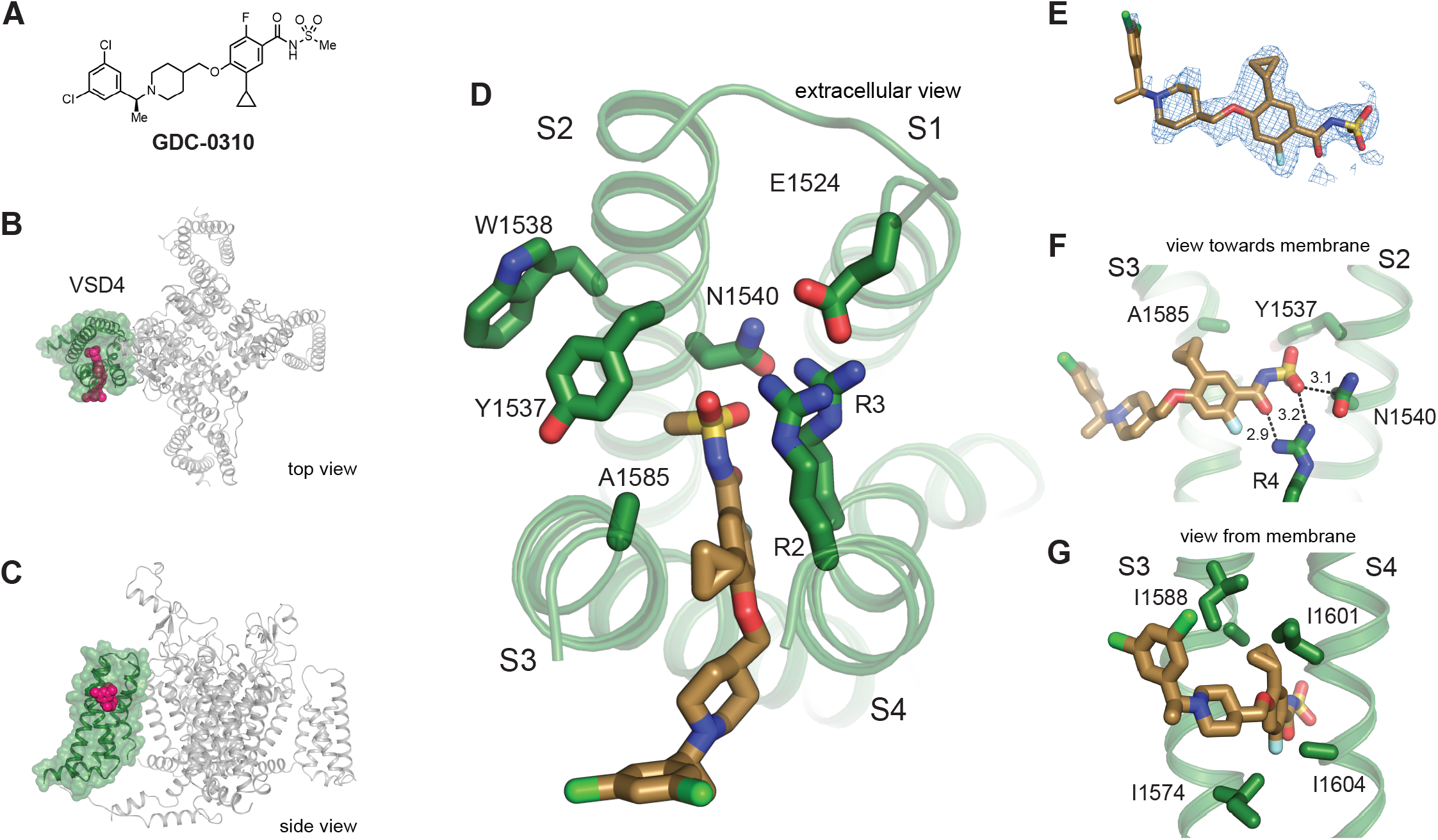
**(A)** Chemical structures of acylsulfonamide **GDC-0310. (B)** Top and Side views of VSD4-NaV1.7-NaVPas channel bound to **GDC-0310**. VSD4 is highlighted in green, **GDC-0310** in magenta. **(D)** Extracellular view of VSD4-NaV1.7-NaVPas acylsulfonamide receptor site is shown with select side chains rendered as sticks. **(E)** The cryoEM map surrounding the ligand **GDC-0310** is shown in mesh representation. **(F)** View towards the membrane highlighting key interactions with the **GDC-0310** warhead. **(G)** View from the membrane highlighting van der Waals interactions with the **GDC-0310** cyclopropyl substituent. The alpha-methylbenzylamine tail region sits almost entirely within the lipid bilayer.

It is notable that the cyclopropyl substituent of **GDC-0310** makes effective van der Waals interactions with I1574 and I1588 from S3 and I1601 and I1604 on S4 (Figure 3G), while the alpha-methylbenzylamine “tail” region sits almost entirely within the lipid bilayer (Figure 3A, C), where density for this portion of the inhibitor is only poorly resolved (Figure 3E). Considering the relative depth of the **GDC-0310** binding site in relation to the membrane-water interface (∼8 Å), our structure suggest a membrane access pathway for acylsulfonamides to the VSD4 receptor site, which has important implications for understanding the pharmacology, SAR, and potential development liabilities of the series.^32^ Notably, this structure also reveals that the Tyr1537 side-chain on S2 exists in a down rotamer and does not contact **GDC-0310** directly, which provides the first direct structural rationale for why the selectivity profiles between the acyl and arylsulfonamide classes differ substantially (Figure 3D, *vide infra*).^33^

### Structure-based rational design of hybrid NaV1.7 inhibitors

Upon inspection of the superimposed **GNE-3565**-aryl and **GDC-0310**-acyl structures, we could immediately envision a novel class of hybrid inhibitors that would simultaneously occupy both the aryl- and acylsulfonamide binding pockets (Figure 4A, B). Such an enlarged hybrid binding pocket would offer unique opportunities to gain potency directly from ligand-protein interactions, which might enable removal of the hydrophobic lipid-facing tail groups seen in all previous generations of VSD4-targeting NaV channel inhibitors. These groups have typically been critical for maintaining potency, but use of the plasma membrane to drive membrane-bound target occupancy is largely nonspecific, and potentially introduces off-target liabilities that can contribute to toxicity. However, it was unclear if this hybrid conformation of the VSD4 domain could accommodate the described inhibitors and/or whether it was accessible during any stage of NaV1.7 channel activation. We therefore set out to demonstrate proof-of-concept that appropriately designed small molecules could induce this hypothesized receptor state while inhibiting channel function with meaningful potency.

**Figure 4:**
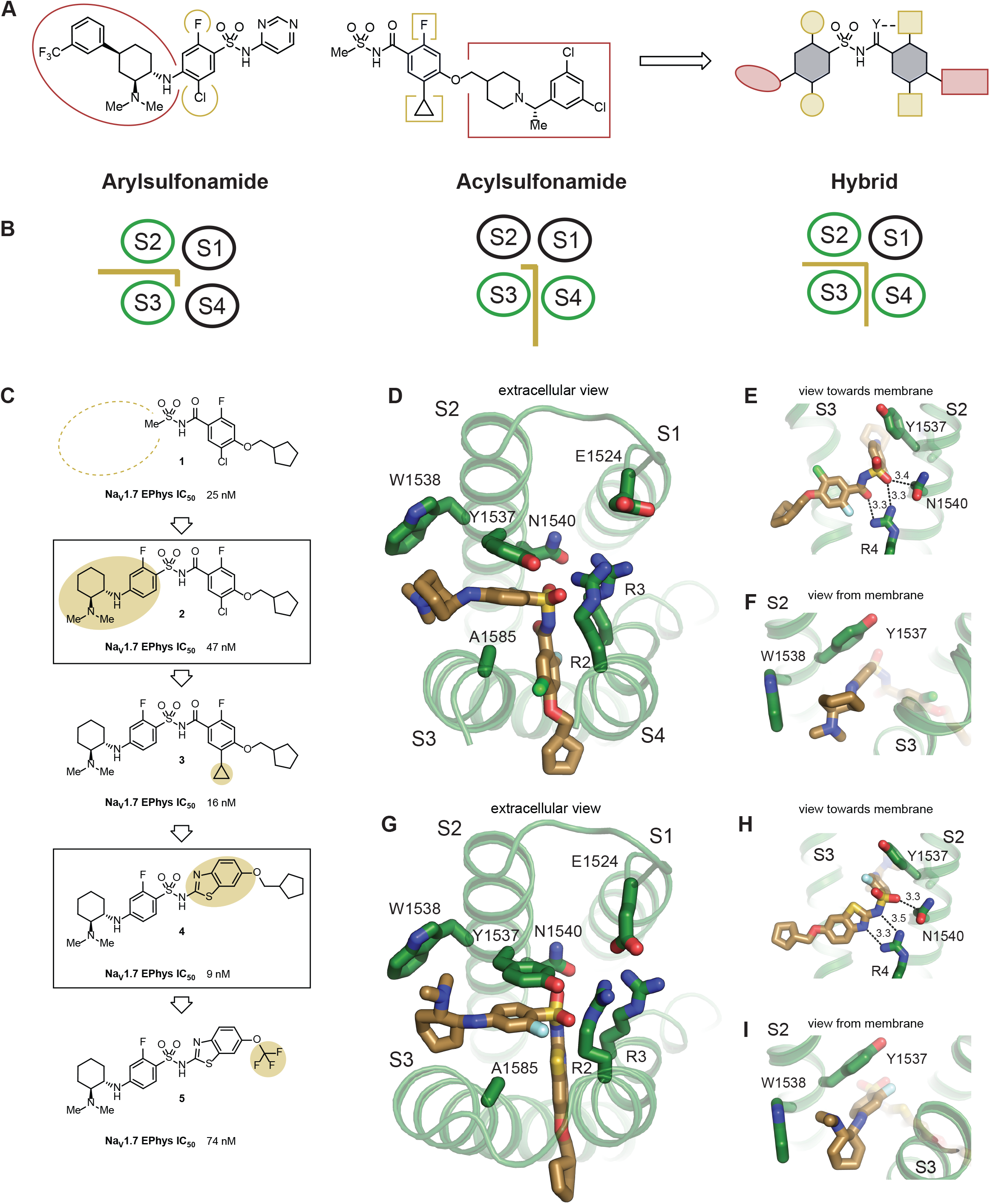
**(A)** Illustration of hybrid molecule design approach. **(B)** Arylsulfonamide, acylsulfonamide and hybrid molecule poses **(C)** Hybridization strategy and molecule optimization. Highlighted are molecule **2** and molecule **4 (D)** Extracellular view of VSD4-NaV1.7-NaVPas bound to the hybrid molecule **2. (E)** View towards the membrane highlighting key interactions with the anionic group. **(F)** View from the membrane highlighting the lack of a stacking interaction between Y1537 and the phenyl ring. **(G)** Extracellular view of VSD4-NaV1.7-NaVPas bound to the hybrid molecule **4. (H)** View towards the membrane highlighting key interactions with the anionic group. **(I)** View from the membrane highlighting the υ-stacking interaction between Y1537 and the phenyl ring.

**GDC-0310** was a relatively unattractive starting point for hybridization because of its high molecular weight (543 g/mol) and lipophilicity (cLogP = 5.2). Accordingly, analysis of historical data identified progenitor compound **1**, which offers reasonable potency against NaV1.7 (IC_50_ = 25 nM, n = 8; Fig 4C) and reduced molecular weight (350 g/mol) and lipophilicity (cLogP = 2.7). Seeking to leverage existing knowledge of the arylsulfonamide binding pocket, molecule **2** was synthesized in an attempt to directly graft on a significant portion of the **GNE-3565** scaffold. Gratifyingly, this molecule retained substantial potency, affording channel blockade at an IC_50_ of 47 nM (n = 8).

A cryo-EM structure of **2** bound to the VSD4 NaV1.7-NaVPas channel confirmed that the molecule adopts the desired hybrid binding mode (Figure 4D). Here, the anionic group of the novel hybrid compound binds in the same position as in the aryl and acyl poses, interacting closely with R4, as seen for the aryl-like binding (Figure 4E). The aryl- and acylsulfonamide-derived structural motifs of **2** occupy their expected positions in the pockets formed between S2/S3 and S3/S4, respectively. Notably, in this hybrid configuration, Y1537 adopts a similar arrangement as observed for the arylsulfonamides (Figure 4F).

Encouraged by this result, we hoped to draw on previously established knowledge of acylsulfonamide SAR to improve the potency of this compound. Established SAR within the acylsulfonamide series suggested that replacement of the chloro substituent on the benzamide fragment of the molecule with a cyclopropane would typically improve potency. This SAR proved translatable to the hybrid class molecules, resulting in inhibitor **3** (IC_50_ = 16 nM, n = 12; Figure 4C).

Further analysis of the cryo-EM structure of **2** revealed that the aryl ring in the S2/S3 binding pocket was shifted ∼1 Å away from the VSD4 core when compared to arylsulfonamides such as **GNE-3565** (data not shown). The aryl ring in the S3/S4 pocket exhibited better overlap with the analogous substituents in the **GDC-0310** structure. We hypothesized that the acylsulfonamide moiety of **2**, which places the two aryl substituents 4.7 Å apart, might not be fully optimized for use in hybrid inhibitors. In comparison, an arylsulfonamide moiety would position these two groups at a distance of ∼3.3 Å. On this basis we designed *N-*benzothiazolyl sulfonamide **4**, which demonstrated potent NaV1.7 inhibition (IC_50_ = 9 nM, n = 8, Figure 4C).

Our last hypothesis was again validated with a cryo-EM co-structure of **4** bound to the VSD4 NaV1.7-NaVPas channel (Figure 4G). Here, we observed the typical face-on pi-stacking orientation of the S2/S3 substituent with Y1537 while maintaining efficient space filling in the S3/S4 pocket. Moreover, the efficient occupation of the non-membrane exposed regions of the binding pocket permitted the removal of the lipophilic tail of the molecule in **5**, where a CF_3_ group replaces the cyclopentylmethyl substituent. Although this molecule shows reduced potency (IC_50_ = 74 nM, n = 8), lack of substantial small molecule contact with the lipid bilayer is an uncommon feature in all other known NaV1.7 VSD4 domain inhibitors. These molecules serve as a key proof of concept, highlighting the power of cryoEM to yield new structural insights that can inspire the design of novel, structurally differentiated small molecule inhibitors.

## Discussion

Iterative structure-based design is among the most powerful techniques to facilitate the development of small molecule drug candidates. Although a variety of technologies are capable of providing information about small molecule ligand-protein interactions, SBDD has traditionally been synonymous with the use of X-ray crystallography, with other techniques failing to rival its high resolution and fast cycle times. Here, we have demonstrated that a robust cryo-EM structure pipeline is sufficient to quickly provide co-structures at resolutions suitable for use in an active medicinal chemistry optimization campaign. Moreover, we have highlighted the power of structural methods, and cryo-EM in particular, to provide new and important insights into inhibitor design through the discovery of a previously unknown binding mode for acylsulfonamide NaV1.7 inhibitors. We further note that our use of an engineered VSD4-NaV1.7-NaVPas chimeric channel construct was essential to produce high yields of recombinant protein capable of routinely returning multiple cryo-EM samples per preparation. This system repeatedly delivered high-resolution depictions of the VSD4 inhibitor binding site and thus strongly contrasts with all prior accounts of cryo-EM structures of full-length human NaV1.7 channels.^22,23^ Importantly, high quality cryo-EM maps at VSD4 in complex with ICA121431 have recently been reported from full-length human NaV1.3 channel protein,^37^ indicating that SBDD efforts targeting a particular NaV channel subtype would benefit from exploring both engineered and native channel sample options at early project stages.

The co-structure of **GDC-0310** was critical to contextualize historical differences between aryl- and acylsulfonamide NaV1.7 inhibitors, which include divergent SAR in the linker phenyl ring and tail regions, disparate NaV family selectivity patterns, and differences in on-rates and off-rates.^12,15,33-36^ Previous efforts to model acylsulfonamide binding have attempted to dock molecules into the **GX-936** co-structure and have proposed subtle changes in molecular register and/or pose to explain the non-translatable SAR.^15,38^ Here, we demonstrate that the molecular features of aryl- and acylsulfonamide inhibitors diverge because they occupy distinct, but overlapping, pockets within the VSD4 domain.

The selectivity determinants of arylsulfonamide NaV1.7 inhibitors have been well studied. Mutation of key non-conserved residues proximal to the binding pocket observed in the **GX-936** co-structure have been shown to alter NaV1.7 potency consistent with selectivity patterns observed for other NaV family members.^14^ Typically, arylsulfonamides have shown high levels of selectivity against NaV1.5, NaV1.1, and NaV1.4, but lower selectivity against NaV1.2 and NaV1.6. In comparison, several acylsulfonamides have been characterized with very high selectivity over NaV1.2 and NaV1.6.^33^ Interestingly, residues forming the contours of the acylsulfonamide binding pocket show high sequence homology across NaV isoforms, suggesting that selectivity for this class is likely an allosteric phenomenon. Nonetheless, the disparate binding pockets for these two inhibitor classes provide a clear rationale for their historically divergent patterns of selectivity.

Our structure of **GDC-0310** provides a structural rationale for the relative slow dissociation kinetics of acylsulfonamide inhibitors compared to arylsulfonamides.^33^ Specifically, the acylsulfonamide pocket is buried more deeply into the plasma membrane than the arylsulfonamide pocket, and may be available only through a membrane-access pathway that first involves partitioning of the molecule into the intramembrane space. In comparison, the arylsulfonamide pocket is open to the outer membrane interface, offering access from the solvent compartment. Slow dissociation kinetics are known to be associated with membrane-access pathways for small molecules binding to transmembrane proteins.^32,39^ For example, the long duration of action of salmeterol (a long-acting β_2_ adrenergic receptor agonist) is proposed to be a result of its membrane partitioning properties. In comparison, salbutamol, which binds to the same target and site but through a solvent-access pathway, shows much faster dissociation from the receptor.^40-46^

In addition to offering valuable information for retrospective analysis of historical NaV1.7 VSD4 domain inhibitors, the structure of **GDC-0310** inspired the development of a class of structurally differentiated molecules that bind to a novel conformation of the receptor. These hybrid molecules bridge both the S2/S3 and S3/S4 helical gaps, with the anionic moiety situated centrally, interacting with the arginine network at the core of the domain. We have demonstrated proof-of-concept that potent hybrid inhibitors can be designed with fewer membrane-associated elements than prior generation NaV1.7 inhibitors, and anticipate that this will offer opportunities to develop molecules with improved lipophilic ligand efficiency and/or differentiated pharmacokinetic profiles. Hybrid inhibitors can also access selectivity determinants from both the S2/S3 and S3/S4 pockets simultaneously, offering a tantalizing opportunity to develop exquisite selectivity over other NaV isoforms.

Beyond the scope of NaV channel inhibitors, our experiences may also offer lessons translatable to small molecule modulators of other transmembrane proteins. Specifically, it is noteworthy to observe that both aryl- and acylsulfonamide inhibitors have access to cryptic pockets that are not observable in the co-structures of the parent proteins. Designing small molecules to “push” on a structural protein element is a common strategy in medicinal chemistry, but it is often challenging to predict when such efforts will afford new, well-defined binding pockets. As such, these designs are often met with failure and quickly abandoned. However, some scenarios exist in which a cryptic pocket is more likely to be found. In these cases, medicinal chemists are more often emboldened to continue experimenting with designs that push into new areas of a protein binding pocket, even when met with initial failure. For example, type II kinase inhibitors can be rationally designed starting from a type I inhibitor structure by appending groups that push through the DFG into the cryptic back pocket.^47^ This pocket is not visible from the type I inhibitor structure, but knowledge of the protein dynamics associated with the DFG-in/DFG-out transition provides a rationale for exploring these designs. We speculate that transmembrane intrahelical domains, particularly those in ion channels, transporters, and GPCRs, may present similarly privileged protein structural motifs that are rich with opportunities for the discovery of druggable cryptic pockets.

In summary, we have developed a robust pipeline for high-resolution structural determination of small-molecule bound NaV1.7 channels suitable for use in iterative structure-based drug design campaigns. We also report the first co-structure of an acylsulfonamide inhibitor bound to the VSD4 domain of NaV1.7, revealing an unexpected and unique binding mode. This finding provides clear rationale for previously unexplainable divergence in the in vitro pharmacological behavior between the two inhibitor classes, and inspired the development of hybrid inhibitors that merge structural features from both inhibitor classes, engaging the receptor in a novel conformation. Our findings highlight the power of cryoEM as an enabling drug discovery technology and offer interesting learnings regarding ion channel structural dynamics that are potentially applicable to other related targets and target classes.

## Materials and Methods

### **Generation of** VSD4-NaV1.7-NaVPas channel constructs

VSD4-NaV1.7-NaVPas chimeric constructs were used as described previously.^21^ In brief, optimized coding DNA for NaV1.7 VSD4-NaVPaS chimeras with N-terminal tandem StrepII and FLAG tag was cloned into a pRK vector with CMV promoter. HEK293 cells in suspension were cultured in SMM 293T-I medium under 5% CO_2_ at 37°C and transfected using PEI when the cell density reached 4 × 10^6^ cells per ml. Transfected cells were harvested 48 hours after transfection. The Dc1a toxin coding DNA from *Diguetia canities* was cloned into a modified pAcGP67A vector downstream of the polyhedron promoter and an N-terminal 6x HIS tag. Recombinant baculovirus was generated using the Baculogold system (BD Biosciences) and *Trichoplusia ni* cells were infected for protein production. The supernatant was harvested 48 h post-infection.

### Protein expression and purification

One hundred fifty grams of cell pellet was resuspended in 500mL of 25mM Hepes pH 7.5, 200mM NaCl, 1ug/mL benzonase, 1mM PMSF, and Roche protease inhibitor tablets. Cells were lysed by dounce homogenization. Proteins were solubilized by addition of 2% GDN (Avanti) with 0.3% cholesteryl hemisuccinate (Avanti) for 2 hours at 4°C. Cell debris was separated by ultracentrifugation at 40,000 rpm at 4°C. Affinity purification using anti-Flag resin was performed by batch binding for 1 hour at 4°C. The resin was washed with 5CV Purification Buffer (25mM Hepes pH 7.5, 200mM NaCl, 0.01% GDN). Another 5CV wash was performed using Purification Buffer supplemented with 5mM ATP and 10mM MgCl_2_. Proteins were eluted in 6 CV Purification Buffer with 300ug/mL Flag peptide. Proteins were subjected to another round of affinity purification using StrepTactin resin (IBA). Proteins were eluted, and concentrated to ∼ 5 mg/mL. For the compound **2** containing sample, instead of nanodisc reconstitution VSD4-NaV1.7-NaVPas was incubated overnight with Dc1a toxin at a 2:1 molar ration of toxin:VSD4-NaV1.7-NaVPas. The eluted sample was concentrated to 100uL, and separated on Superose 6 3.2/300.

For nanodisc reconstitution used for the samples containing **GNE-3565, GDC-0310** and compound **4**, a 200-molar excess of lipid mix (3POPC:1POPE:1POPG resuspended in 50mM Hepes pH7.5, 100mM NaCl, 5mM MgCl2, 1% CHAPS) was added to detergent-solubilized protein and incubated on ice for 30 minutes. A four-molar excess of scaffold protein (MSP1E3D1, Sigma) was added to the protein-lipid mix, and incubated on ice for another 30 minutes. To remove detergent, BioBeads (Bio-Rad) were added to 0.25 mg/mL, and incubated overnight at 4°C. To remove the empty nanodiscs, the sample was subjected to a round of affinity purification using Strep-Tactin (IBA). The eluted sample was concentrated to 100uL, and separated on Superose 6 3.2/300. Peak fractions were combined, and split into four samples. Fifty uM of the small molecule of interest was added to each sample, and incubated at 22°C for 10 minutes. The nanodisc reconstituted samples were crosslinked with 0.05% glutaraldehyde (Electron Microscopy Sciences) then quenched with 1M Tris pH 7.0. Samples at 2 mg/mL were used for grid freezing.

### CryoEM sample preparation and data acquisition

CryoEM grids for the small molecule VSD4-NaV1.7-NaVPas chimeric complexes shown in this study were prepared as follows: For the **GDC-0310** and compound **4** VSD4-NaV1.7-NaVPas complexes, holey carbon grids (Ultrafoil 25 nM Au R 0.6/1 300 mesh; Quantifoil) were incubated with a thiol reactive, self-assembling reaction mixture of 4mM monothiolalkane(C11)PEG6-OH (11-mercaptoundecyl) hexaethyleneglycol (SPT-0011P6, SensoPath Technologies, Inc., Bozeman, MT). Grids were incubated with this self-assembled monolayer (SAM) solution for 24 hours and afterwards rinsed with EtOH.

3 μl of the sample was applied to the grid and blotted with Vitrobot Mark IV (Thermofisher) using 3.5 s blotting time with 100% humidity and plunge-frozen in liquid ethane cooled by liquid nitrogen. The **GNE-3565** VSD4-NaV1.7-NaVPas complex was, similarly as described above, applied to a holey carbon grid (Ultrafoil 25 nM Au R 1.2/1,3 300 mesh; Quantifoil) pretreated with SAM solution. The grid was blotted single-sided with a Leica EM GP (Leica) using 3 s blotting time with 100% humidity and plunge-frozen in liquid ethane cooled by liquid nitrogen. For compound **2** VSD4-NaV1.7-NaVPas-DC1a complex, holey carbon grids (Ultrafoil 25 nM Au R 2/2 200 mesh; Quantifoil) were glow-discharged for 10 s using the Solarus plasma cleaner (Gatan). 3 μl of the sample was applied to the grid and blotted with Vitrobot Mark IV (Thermofisher) using 2.5 s blotting time with 100% humidity and plunge-frozen in liquid ethane cooled by liquid nitrogen.

Movie stacks for compound **2** VSD4-NaV1.7-NaVPas-DC1a complex were collected using SerialEM^48^ on a Titan Krios operated at 300 keV with bioquantum energy filter equipped with a K2 Summit direct electron detector camera (Gatan). Images were recorded at 165,000 × magnification corresponding to 0.824 Å per pixel, using a 20 eV energy slit. Each image stack contains 50 frames recorded every 0.2 s for an accumulated dose of ∼50 e Å−2 and a total exposure time of 10 s. Images were recorded with a set defocus range of to 1.5 μm.

Movie stacks for **GNE-3565, GDC-0310** and compound **4** VSD4-NaV1.7-NaVPas complexes were collected using SerialEM on a Titan Krios G3i (ThermoFisher Scientific, Waltham, MA) operated at 300 keV with bioquantum energy filter equipped with a K3 Summit direct electron detector camera (Gatan Inc, Pleasanton, CA). Images were recorded in EFTEM mode at 105,000 × magnification corresponding to 0.838 Å per pixel, using a 20 eV energy slit. Each image stack contains 60 frames recorded every 0.05 s for an accumulated dose of ∼60 e Å−2 and a total exposure time of 3 s. Images were recorded with a set defocus range of 0.5 to 1.5 μm.

### CryoEM data processing

CryoEM data were processed using a combination of the RELION^49^ and cisTEM^50^ software packages, similarly as described previously^51^ and as illustrated in Suppl. Figure 2 and 3.

Movies were corrected for frame motion using the MotionCor2^52^ implementation in RELION and contrast-transfer function parameters were fit using the 30-4.5 Å band of the spectrum with CTFFIND-4.^53^ CTF fitted images were filtered on the basis of the detected fit resolution better than 6-10 Å. Particles were picked by template-matching with gautomatch (https://www2.mrc-lmb.cam.ac.uk/research/locally-developed-software/zhang-software/#gauto) using a 30 Å low-pass filtered apo VSD4-NaV1.7-NaVPas reference structure. Particles were sorted during RELION 2D classification and selected particles were imported into cisTEM for 3D refinements. 3D reconstructions were obtained after auto-refine and manual refinements with a mask around the channel (excluding the detergent micelle) and by applying low-pass filter outside the mask (filter resolution 20 Å) and a score threshold of 0.10-0.30, so that only the best-scoring 10-30% of particle images would be included in the 3D reconstruction at each cycle. The weight outside of the mask was set to 0.8. No data beyond 3.0 Å for compound **4**, 3.4 Å for **GDC-0310**, 3.7Å **GNE-3565** and 4.0 Å for compound **2** were used in the refinements. Phenix ResolveCryoEM^54^ density modification was applied to each of the reconstructions to obtained the final map used for model building. Local resolution was determined in cisTEM using a reimplementation of the blocres algorithm.

### Model building and structure analysis

The previously determined VSD4-NaV1.7-NaVPas model (PDB: 6NT3)^21^ was fit as a rigid body into the cryoEM map. After manual adjustments, multiple rounds of real space refinement using the phenix.real_space_refinement tool^55^ was used to correct structural differences between the initial model and the map. The small molecule coordinates were generated with eLBOW,^56^ placed and manually adjusted in Coot,^57^ energy minimized with MOE (Chemical Computing Group ULC,) and refined with real space refinements in Phenix^55^. The model was validated using phenix.validation_cryoem^58^ with built-in MolProbity scoring.^59^ Figures were made using PyMOL (The PyMOL Molecular Graphics System, v.2.07 Schrödinger, LLC), UCSF Chimera,^60^ and UCSF ChimeraX.^61^

### Chemical synthesis

The synthesis of **GNE-3565, GDC-0310**, and compound **1** have been previously reported.^12,15,62^ The synthesis and characterization of compounds **2-5** are described in the supporting information.

### NaV1.7 Syncropatch electrophysiology potency

cDNAs for NaV1.7 (NM_002977) were stably expressed in Chinese Hamster Ovary (CHO) cells. Sodium currents were measured in the whole-cell configuration using Syncropatch 384PE (NanIon Technologies, Germany). 1NPC®-384 chips with custom medium resistance and single hole mode were used. Internal solution consisted of (in mM): 110 CsCl, 10 CsCl, 20 EGTA, and 10 Hepes (pH adjusted to 7.2); and external solution contains (in mM): 60 NMDG, 80 NaCl, 4 KCl, 1 MgCl_2_, 2 CaCl_2_, 2 D-Glucose monohydrate, 10 Hepes (pH adjusted to 7.4 with NaOH).

After system flushing, testing compounds were dissolved in external solution containing 0.1% Pluronic F-127. 10µl cells were added to the chip from a cell hotel, and a negative pressure of −50 mBar was applied to form a seal. Following treatment with seal enhancer solution and wash-off with external solution, negative pressure of −250 mbar was applied for 1 second to achieve the whole-cell configuration, followed by three washing steps in external solution. 20µl of compounds were added to 40µl in each well (1:3 dilution of compounds), and after mixing, 20µl was removed so the volume was retained at 40 ul. After approximately 13 minutes recording, 20µl/well of 2 uM TTX was added to achieve full block.

For voltage protocol, a holding potential of -50 mV was applied during the whole experiment. A depolarizing step was applied to -10mV for 10ms, followed by a hyperpolarization step to -150 mV for 20 ms to allow channel recovery from inactivation. A second depolarizing step was applied from -150mV to -10mV for 10 ms, where currents were measured to derive blocking effects of compounds. Inhibition was determined based on 7.5 min of compound incubation.

## Acknowledgments

We are grateful to BMR and RMG for construct generation and protein scale-up. We are thankful to Cameron Noland for help with the biochemistry of the VSD4-NaV1.7-NaVPas channel system. We thank Tommy Lai and Wenfeng Liu for chemistry support.

