## Supplementary Information for "CryoEM reveals unprecedented binding site for Na_V_1.7 inhibitors enabling rational design of potent hybrid inhibitors"

**Table S1. Cryo-EM data collection, refinement and validation statistics.**

| | **GNE-3565** VSD4-$Na_V$1.7-$Na_V$Pas complex | **GDC-0310** VSD4-$Na_V$1.7-$Na_V$Pas complex | **GNE-1305** VSD4-$Na_V$1.7-$Na_V$Pas complex | **GNE-9296** VSD4-$Na_V$1.7-$Na_V$Pas-DC1a complex |
|---|---|---|---|---|
| **Data Collection** | | | | |
| Magnification | 105,000x | 105,000x | 165,000x | 165,000x |
| Voltage (kV) | 300 | 300 | 300 | 300 |
| Electron exposure (e/Å$^2$) | 64 | 64 | 64 | 48 |
| Defocus range (μm) | 0.5-1.5 | 0.5-1.5 | 0.5-1.5 | 0.5-1.5 |
| Pixel size (Å) | 0.838 | 0.838 | 0.838 | 0.824 |
| Symmetry imposed | C1 | C1 | C1 | C1 |
| Initial particle images | 3,686,837 | 2,253.983 | 2,741,402 | 1,774,062 |
| Final particle images | 964,572 | 795,792 | 1,201,168 | 826,525 |
| Map resolution (Å) overall | 2.9 | 2.5 | 2.2 | 3.1 |
| FSC threshold | 0.143 | 0.143 | 0.143 | 0.143 |
| Map resolution range (Å) | 2.7-33.5 | 2.3-33.5 | 2.1-33.5 | 3.3-33.0 |
| **Refinement** | | | | |
| Initial models used (PDB code) | 6NT3 | 6NT3 | 6NT3 | 6NT3 |
| Model resolution (Å) | 2.9 | 1.9 | 1.7 | 2.5 |
| FSC threshold | 0.5 | 0.5 | 0.5 | 0.5 |
| Model composition | | | | |
| Non-hydrogen atoms | 9396 | 9559 | 9617 | 9734 |
| Protein residues | 1115 | 1115 | 1117 | 1159 |
| Waters | 0 | 80 | 112 | 0 |
| Ligands | 13 | 13 | 14 | 13 |
| B factors (Å$^2$) | | | | |
| Protein | 25.62 | 12.42 | 6.27 | 24.05 |
| Ligand | 35.81 | 15.79 | 10.33 | 40.62 |
| Water | N/A | 8.07 | 6.57 | N/A |
| R.m.s. deviations | | | | |
| Bond lengths (Å) | 0.004 | 0.005 | 0.005 | 0.005 |
| Bond angles (°) | 0.931 | 1.018 | 0.993 | 0.940 |
| Validation | | | | |
| MolProbity score | 1.27 | 1.18 | 1.13 | 1.37 |
| Clashscore | 5.15 | 3.99 | 3.40 | 6.32 |
| Poor rotamers (%) | 0.20 | 0.30 | 0.61 | 0.00 |
| Ramachandran Plot | | | | |
| Favored (%) | 98.45 | 98.27 | 98.19 | 97.90 |
| Allowed (%) | 1.55 | 1.73 | 1.81 | 2.10 |
| Disallowed (%) | 0.00 | 0.00 | 0.00 | 0.00 |

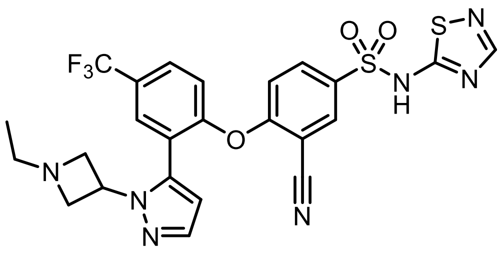

**GX-936**

**Supplementary Figure 1:** Structure of arylsulfonamide inhibitor **GX-936**

**Chemical compound synthesis and characterization**

**General Considerations**

All commercial solvents and reagents were used without additional purification unless indicated otherwise. $^1$H NMR spectra were measured on Bruker Avance III 300, 400, or 500 MHz spectrometers. Chemical shifts (in ppm) were referenced to tetramethylsilane as an internal standard (δ = 0 ppm). Reaction progress was monitored by either a Shimadzu LCMS/UV system with an LC-30 AD solvent pump, Sil-30 AC autosampler, 2020 MS, SPDM30A UV detector, and CTO-20A column oven, using 2−98% acetonitrile/0.1% formic acid (or 0.01% ammonia) over 2.5 min or a Waters Acquity LCMS system using 2−98% acetonitrile/0.1% formic acid (or 0.1% ammonia) over 2 min. Flash column chromatography purifications were performed using a Teledyne Isco Combiflash Rf and Silicycle HP columns. Reverse-phase purification was done on a Phenomenex Gemini-NX C18 (30 × 100 mm, 5 μm) with a gradient of 5−95% acetonitrile/water (with 0.1% NH4OH or 0.1% formic acid) at 60 mL/min over 10 min. Preparative SFC separations were carried out on a PIC Solutions instrument, with conditions specified in the Experimental Section. High-resolution mass spectrometry (HRMS) of final compounds was obtained on a Thermo UHPLC/ QE with a Thermo-Q Exactive mass spectrometry detector using ESI ionization, following elution on an Acquity BEH C18 stationary phase (2.1 mm × 50 mm; 1.7 μm particle size) using a gradient of water/ acetonitrile (3−97% over 7 min with 0.1% formic acid in both phases). Unless stated otherwise, analytical purity was >95% as determined by LCMS using UV 254 nm detection

**Compound 2**

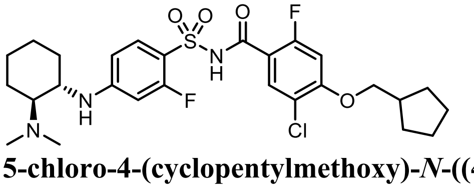

**5-chloro-4-(cyclopentylmethoxy)-*N*-((4-(((1*S*,2*S*)-2-(dimethylamino)cyclohexyl)amino)-2-fluorophenyl)sulfonyl)-2-fluorobenzamide**

**Step 1:**

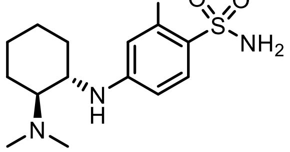

**4-(((1*S*,2*S*)-2-(dimethylamino)cyclohexyl)amino)-2-fluorobenzenesulfonamide**

A solution of (1*S*,2*S*)-*N*$^1$,*N*$^1$-dimethylcyclohexane-1,2-diamine (162 mg, 1.1 mmol), 2,4-difluorobenzenesulfonamide (200 mg, 1.0 mmol) and DIPEA (0.3 mL, 1.7 mmol) in DMSO (3 mL) was stirred at

80 ºC for 16 h. After cooling to room temperature, the reaction was diluted with EtOAc (100 mL), and washed with brine (50 mL x 5). The organic lawyer was dried over anhydrous $Na_2SO_4$, filtered and concentrated in vacuo. The crude residue was purified by silica gel chromatography (solvent gradient: 0 - 10% EtOAc in petroleum ether) to afford the title compound (60 mg, 0.19 mmol) as yellow oil.
LCMS (ESI) m/z: 316.1 $[M+H]^+$.

**Step 2:**

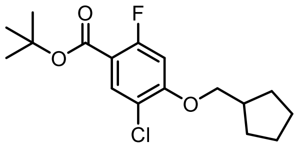

**tert-butyl 5-chloro-4-(cyclopentylmethoxy)-2-fluorobenzoate**

A mixture of tert-butyl 5-chloro-2,4-difluorobenzoate (548 g, 2.20 mol), cyclopentylmethanol (200 g, 2.00 mol) and $Cs_2CO_3$ (1.31 kg, 4.00 mol) in DMSO (3.28 L) was stirred at 80 °C for 5 hrs. The mixture was filtered and filtrate was diluted with EtOAc (3.00 L), washed with water (3.00 L x 3) and brine (1000 mL). The resultant organic layer was dried over $Na_2SO_4$, filtered, concentrated in vacuo. The residue was purified by column chromatography ($SiO_2$, Petroleum ether/Ethyl acetate = 100/1 to 50/1). The title compound (0.50 kg, crude) was obtained as light yellow oil.

**Step 3:**

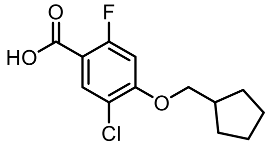

**5-chloro-4-(cyclopentylmethoxy)-2-fluorobenzoic acid**

To a solution of *tert*-butyl 5-chloro-4-(cyclopentylmethoxy)-2-fluorobenzoate (327 g, 994 mmol) in DCM (654 mL) was added TFA (654 mL) at 25 °C. The mixture was stirred at 25 °C for 24 hrs. The reaction mixture was filtered and concentrated under reduced pressure to give a residue that was triturated with 1/1 MTBE/Petroleum ether at 25 °C for 30 mins. The title compound was obtained by filtration as a white solid (83.7 g, 30.5% yield).
$^1$H NMR (400 MHz, $CDCl_3$) $\delta$ 8.04 (d, $J$ = 7.6 Hz, 1H), 6.70 (d, $J$ = 12.4 Hz, 1H), 3.96 (d, $J$ = 6.8 Hz, 2H), 2.42-2.50 (m 1H), 1.86-1.93 (m, 2H), 1.75-1.58 (m, 4H), 1.48-1.35 (m, 2H).

**Step 4:**

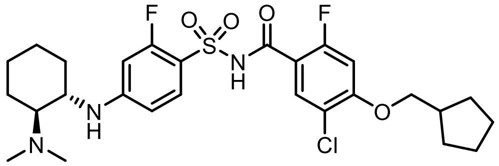

**5-chloro-4-(cyclopentylmethoxy)-*N*-((4-(((1*S*,2*S*)-2-(dimethylamino)cyclohexyl)amino)-2-fluorophenyl)sulfonyl)-2-fluorobenzamide**

To a solution of 4-(((1*S*,2*S*)-2-(dimethylamino)cyclohexyl)amino)-2-fluorobenzenesulfonamide (80 mg, 0.24 mmol) and DMAP (59 mg, 0.48 mmol) in DCM (2 mL) was added EDCI (51 mg, 0.26 mmol) and 5-chloro-4-(cyclopentylmethoxy)-2-fluoro-benzoic acid (72 mg, 0.26 mmol). The reaction was stirred at room temperature for 2 h. The reaction was quenched with 10% aqueous citric acid (5 mL). The reaction was diluted with water (10 mL) and extracted with DCM (10 mL x 3). The combined organic layers were dried over anhydrous $Na_2SO_4$, filtered and concentrated in vacuo. The crude residue was purified by silica gel chromatography (solvent gradient: 0 - 5% MeOH in DCM) to afford the title compound (40 mg, 28%) as a white solid.

$^1$H NMR (400 MHz, DMSO-$d_6$) δ 7.75 (d, $J$ = 7.6 Hz, 1H), 7.55 - 7.48 (m, 1H), 6.94 (d, $J$ = 12.4 Hz, 1H), 6.50 (s, 1H), 6.47 (s, 1H), 6.07 (d, $J$ = 10.0 Hz, 1H), 3.94 (d, $J$ = 6.8 Hz, 2H), 3.79 - 3.64 (m, 1H), 3.15 - 3.05 (m, 1H), 2.66 (s, 6H), 2.35 - 2.25 (m, 1H), 2.09 - 1.94 (m, 2H), 1.84 - 1.69 (m, 3H), 1.63 - 1.14 (m, 11H).
HRMS *m/z* calcd for $C_{27}H_{34}ClF_2N_3O_4S$ $[M+H]^+$: 570.1999. Found 570.2003.

**Compound 3**

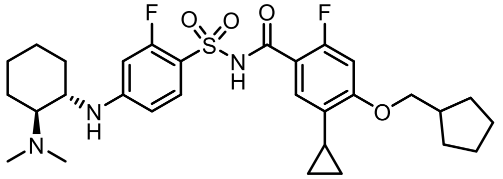

**4-(cyclopentylmethoxy)-5-cyclopropyl-N-((4-(((1S,2S)-2-(dimethylamino)cyclohexyl)amino)-2-fluorophenyl)sulfonyl)-2-fluorobenzamide**

Following the procedure described in compound **3** and replacing 5-chloro-4-(cyclopentylmethoxy)-2-fluorobenzoic acid with 4-(cyclopentylmethoxy)-5-cyclopropyl-2-fluorobenzoic acid, the title compound was obtained as a white solid.
$^1$H NMR (400 MHz, DMSO-$d_6$) δ 7.50 (t, $J$ = 8.8 Hz, 1H), 7.20 (d, $J$ = 8.8 Hz, 1H), 6.65 (d, $J$ = 12.0 Hz, 1H), 6.52 - 6.34 (m, 2H), 5.99 (s, 1H), 3.87 (d, $J$ = 6.8 Hz, 2H), 3.72 - 3.52 (m, 1H), 3.05 - 2.81 (m, 1H), 2.61 - 2.52 (m, 6H), 2.37 - 2.26 (m, 1H), 2.07 - 1.90 (m, 3H), 1.87 - 1.71 (m, 3H), 1.66 - 1.52 (m, 5H), 1.45 - 1.30 (m, 4H), 1.29 - 1.20 (m, 1H), 1.15 - 1.05 (m, 1H), 0.91 - 0.80 (m, 2H), 0.60 - 0.45 (m, 2H).
HRMS *m/z* calcd for $C_{30}H_{39}F_2N_3O_4S$ $[M+H]^+$: 576.2702. Found 576.2713.

**Compound 4**

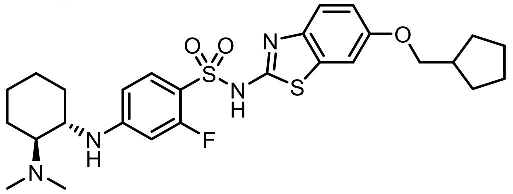

**N-(6-(cyclopentylmethoxy)benzo[d]thiazol-2-yl)-4-(((1S,2S)-2-(dimethylamino)cyclohexyl)amino)-2-fluorobenzenesulfonamide**

**Step 1:**

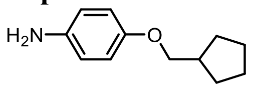

**4-(cyclopentylmethoxy)aniline**

To a stirred solution of NaH (1.32 g, 54.98 mmol, 60% in mineral oil) in DMF (20 mL) was added 4-aminophenol (2 g, 18.33 mmol) at 0 ºC under nitrogen atmosphere. After stirring at 0 ºC for 10 min, (bromomethyl)cyclopentane (4.48 g, 27.49 mmol) was added at 0 ºC. The reaction was stirred at room temperature for 16 h. The reaction was quenched with water (100 mL) and extracted with EtOAc (100 mL x 3). The combined organic lawyers were washed with brine (100 mL), dried over anhydrous $Na_2SO_4$, filtered and concentrated in vacuo. The crude residue was purified by silica gel chromatography (solvent gradient: 10 - 20% EtOAc in petroleum ether) to afford the title compound (1.35 g, 39%) as black oil.
$^1$H NMR (400 MHz, $CDCl_3$) δ 6.77 - 6.74 (m, 2H), 6.66 - 6.63 (m, 2H), 3.76 (d, $J$ = 7.2 Hz, 2H), 3.42 (s, 2H), 2.38 - 2.27 (m, 1H), 1.86 - 1.78 (m, 2H), 1.64 - 1.58 (m, 4H), 1.39 - 1.30 (m, 2H).
LCMS (ESI) m/z: 192.2 $[M+H]^+$.

**Step 2:**

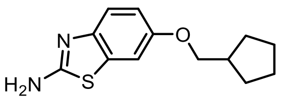

**6-(cyclopentylmethoxy)benzo[*d*]thiazol-2-amine**

The solution of 4-(cyclopentylmethoxy)aniline (1.30 g, 7.06 mmol) and potassium thiocyanate (685 mg, 7.06 mmol) in acetic acid (7.5 mL) was stirred at 0 °C for 20 min. Bromine (0.36 mL, 7.06 mmol) in acetic acid (3.5 mL) was added slowly and keep the temperature below 10 °C. Then, the mixture was stirred at room temperature for 18 h. The reaction was filtered and the filter cake was washed with acetic acid (5 mL). The filtrate was concentrated in vacuo and the crude reside was diluted with hot water (5 mL) and basified to pH>11 with $NH_3$•$H_2O$. The resulting precipitate was filtered and the filter cake was washed with water (5 mL). The filter cake was diluted with DCM (20 mL), dried over anhydrous $Na_2SO_4$, filtered and concentrated in vacuo. The crude residue was purified by silica gel chromatography (solvent gradient: 0 - 14% EtOAc in petroleum ether) to afford the title compound (800 mg, 46%) as a gray solid.

$^1$H NMR (400 MHz, $CDCl_3$) δ 7.43 (d, *J* = 8.8 Hz, 1H), 7.13 (d, *J* = 6.0 Hz, 1H), 6.93 - 6.90 (m, 1H), 5.22 (s, 2H), 3.84 (d, *J* = 7.2 Hz, 2H), 2.41 - 2.33 (m, 1H), 1.89 - 1.81 (m, 2H), 1.68 - 1.58 (m, 4H), 1.41 - 1.33 (m, 2H).

LCMS (ESI) m/z: 249.0 $[M+H]^+$.

**Step 3:**

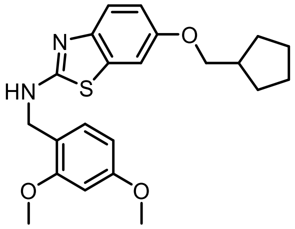

**6-(cyclopentylmethoxy)-N-(2,4-dimethoxybenzyl)benzo[d]thiazol-2-amine**

To a stirred solution of 6-(cyclopentylmethoxy)benzo[d]thiazol-2-amine (0.71 g, 2.65 mmol) and 2,4-dimethoxybenzaldehyde (0.4 g, 2.41 mmol) in DCM (12 mL) was added TiCl(O*i*-Pr)$_3$ (1.86 mL, 5.54 mmol) in one portion under nitrogen atmosphere. The solution was stirred for 10 min before the portion wise addition of $NaBH(OAc)_3$ (1.53 g, 7.22 mmol) at 0 °C. The reaction was stirred at room temperature for 16 h. The reaction was quenched with saturated aqueous $NaHCO_3$ solution (50 mL), extracted with DCM (50 mL x 3). The combined organic lawyers were washed with brine (50 mL), dried over anhydrous $Na_2SO_4$, filtered and concentrated in vacuo. The crude residue was purified by silica gel chromatography (solvent gradient: 0 - 25% EtOAc in petroleum ether) to afford the title compound (0.53 g, 53%) as a white solid.

**Step 4:**

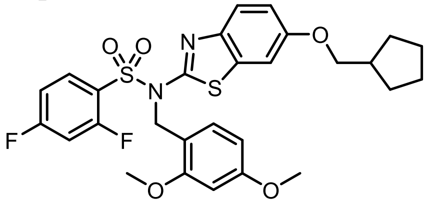

**N-(6-(cyclopentylmethoxy)benzo[d]thiazol-2-yl)-N-(2,4-dimethoxybenzyl)-2,4-difluorobenzenesulfonamide**

To a solution of 6-(cyclopentylmethoxy)-N-(2,4-dimethoxybenzyl)benzo[d]thiazol-2-amine (170 mg, 0.52 mmol) in THF (2 mL) was added LiHMDS (0.62 mL, 0.62 mmol, 1 M) at -78 °C. The reaction was stirred for 30 min at 0 °C and a solution of 2,4-difluorobenzenesulfonylchloride (0.22 g, 1.03 mmol) in THF (2 mL) was added dropwise at -78 °C. After the addition was complete, the cooling bath was removed. The reaction mixture was stirred at room temperature for 3 h. The reaction was diluted with water (30 mL) and extracted with EtOAc (50 mL x 3). The combined organic lawyers were washed with brine (50 mL), dried over anhydrous $Na_2SO_4$, filtered and concentrated in vacuo. The crude residue was purified by silica gel chromatography (solvent gradient: 0 - 30% EtOAc in petroleum ether) to afford the title compound (160 mg, 52%) as a white solid.

**Step 5:**

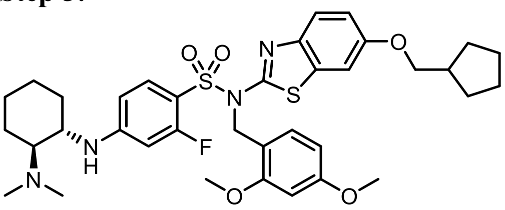

**N-(6-(cyclopentylmethoxy)benzo[d]thiazol-2-yl)-N-(2,4-dimethoxybenzyl)-4-(((1S,2S)-2-(dimethylamino)cyclohexyl)amino)-2-fluorobenzenesulfonamide**

To a solution of *N*-(6-bromothiazolo[4,5-*b*]pyridin-2-yl)-4-(((1*S*,2*S*)-2-(dimethylamino)cyclohexyl)amino)-2-fluorobenzenesulfonamide (40 mg, 0.07 mmol) in DMSO (1 mL) was added DIPEA (14 mg, 0.11 mmol) and (1*S*,2*S*)-$N_1$,$N_1$-dimethylcyclohexane-1,2-diamine (15 mg, 0.11 mmol) at room temperature. The reaction mixture was stirred at room temperature for 20 h. The reaction was quenched with saturated aqueous $NH_4Cl$ (20 mL), extracted with EtOAc (30 mL x 3). The combined organic layers were washed with brine (30 mL), dried over anhydrous $Na_2SO_4$, filtered and concentrated in vacuo to afford the title compound (48 mg, crude) as yellow oil that required no further purification.

**Step 6:**

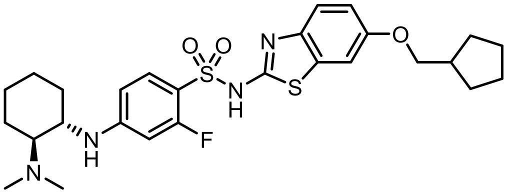

**N-(6-(cyclopentylmethoxy)benzo[d]thiazol-2-yl)-4-(((1S,2S)-2-(dimethylamino)cyclohexyl)amino)-2-fluorobenzenesulfonamide**

A solution of *N*-(6-(cyclopentylmethoxy)benzo[d]thiazol-2-yl)-*N*-(2,4-dimethoxybenzyl)-4-(((1*S*,2*S*)-2-(dimethylamino)cyclohexyl)amino)-2-fluorobenzenesulfonamide (60 mg, 0.08 mmol) in HCOOH (3 mL) was stirred at room temperature for 16 h. The mixture was concentrated in vacuo and the crude residue was purified by reverse phase chromatography (acetonitrile 35 - 65% / 0.2% HCOOH in water) to afford the title compound (2 mg, 4%) as a white solid

$^1$H NMR (400 MHz, DMSO-$d_6$) δ 7.49 (t, *J* = 8.8 Hz, 1H), 7.26 (d, *J* = 2.4 Hz, 1H), 7.16 (d, *J* = 8.8 Hz, 1H), 6.84 - 6.81 (m, 1H), 6.42 - 6.36 (m, 3H), 3.79 (d, *J* = 6.8 Hz, 2H), 3.47 - 3.40 (m, 1H), 2.82 - 2.75 (m, 1H), 2.42 (s, 6H), 2.33 - 2.24 (m, 1H), 2.02 - 1.92 (m, 2H), 1.79 - 1.71 (m, 3H), 1.61 - 1.48 (m, 5H), 1.34 - 1.20 (m, 6H).

HRMS (ESI) m/z: 547.2196 $[M+H]^+$

**Compound 5**

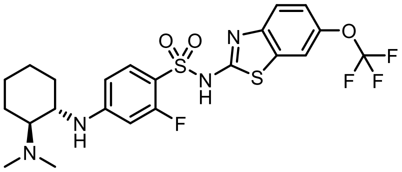

**4-(((1S,2S)-2-(dimethylamino)cyclohexyl)amino)-2-fluoro-N-(6-(trifluoromethoxy)benzo[d]thiazol-2-yl)benzenesulfonamide**

Following the procedure described in compound **6** and replacing 4-(cyclopentylmethoxy)aniline with 4-(trifluoromethoxy)aniline, the title compound was obtained as a white solid.

$^1$H NMR (400 MHz, DMSO-$d_6$) δ 7.64 (m, 1H), 7.52 (m, 1H), 7.30 (d, *J* = 8.8 Hz, 1H), 7.11 (m, 1H), 6.44-6.37 (m, 3 H), 3.68 (m, 1H), 3.06 (m, 1H), 2.60 (s, 6 H), 2.00 (m, 2H), 1.79 (m, 1H), 1.61 (m, 1H), 1.40- 1.14 (m, 4 H).

HRMS (ESI) m/z: 533.1286 $[M+H]^+$.

**Supplementary Figure Legends:**

**Supplementary Figure 2: (A)** VSD4-Na$_V$1.7-Na$_V$Pas channel protein expression and purification scheme **(B)** Example size exclusion chromatogram and SDS-PAGE of nanodisc-reconstituted VSD4-Na$_V$1.7-Na$_V$Pas channel sample.

**Supplementary Figure 3: (A)** Example cryo-EM micrograph image of the VSD4-Na$_V$1.7-Na$_V$Pas channel. **(B)** Representative 2D-class averages of selected particles. **(C)** Data collection and processing workflow for VSD4-Na$_V$1.7-Na$_V$Pas channel bound to compound **4**. **(D)** Heat map representation of the distribution of assigned particle orientations. **(E)** FSC between two half datasets yields a global resolution estimate of approximately 2.2 Å resolution from the refinement using an overall mask of the VSD4-Na$_V$1.7-Na$_V$Pas channel bound to compound **4** **(F)** Local resolution of maps. **(G)** Data collection and processing workflow for VSD4-Na$_V$1.7-Na$_V$Pas channel bound to **GDC-0310**. **(H)** Heat map representation of the distribution of assigned particle orientations. **(I)** FSC between two half datasets yields a global resolution estimate of approximately 2.5 Å resolution from the refinement using an overall mask of the VSD4-Na$_V$1.7-Na$_V$Pas channel bound to **GDC-0310**. **(J)** Local resolution of maps.

**Supplementary Figure 4: (A)** Data collection and processing workflow for VSD4-Na$_V$1.7-Na$_V$Pas channel bound to compound **2**. **(B)** Heat map representation of the distribution of assigned particle orientations. **(C)** FSC between two half datasets yields a global resolution estimate of approximately 2.9 Å resolution from the refinement using an overall mask of the VSD4-Na$_V$1.7-Na$_V$Pas channel bound to compound **2** **(D)** Local resolution of maps. **(E)** Data collection and processing workflow for VSD4-Na$_V$1.7-Na$_V$Pas channel bound to compound **2**. **(F)** Heat map representation of the distribution of assigned particle orientations. **(G)** FSC between two half datasets yields a global resolution estimate of approximately 2.5 Å resolution from the refinement using an overall mask of the VSD4-Na$_V$1.7-Na$_V$Pas channel bound to compound **2**. **(H)** Local resolution of maps.

**Supplementary Figure 5: (A)** The cryoEM map surrounding the ligand compound **2** is shown in mesh representation. **(B)** The cryoEM map surrounding the ligand compound **4** is shown in mesh representation.

HPLC chromatogram of compound **2**

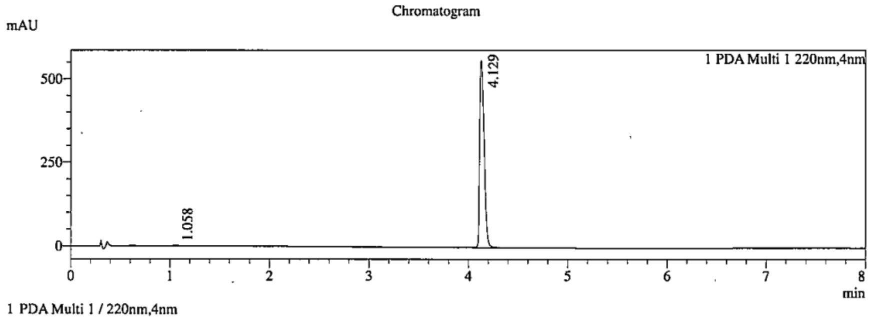

Integration Result

Peak Table

PDA Ch1 220nm

| Peak# | Ret. Time | Height | Height% | USP Width | Area | Area% |
|---|---|---|---|---|---|---|
| 1 | 1.058 | 2105 | 0.384 | 0.071 | 6000 | 0.342 |
| 2 | 4.129 | 546761 | 99.616 | 0.084 | 1750524 | 99.658 |

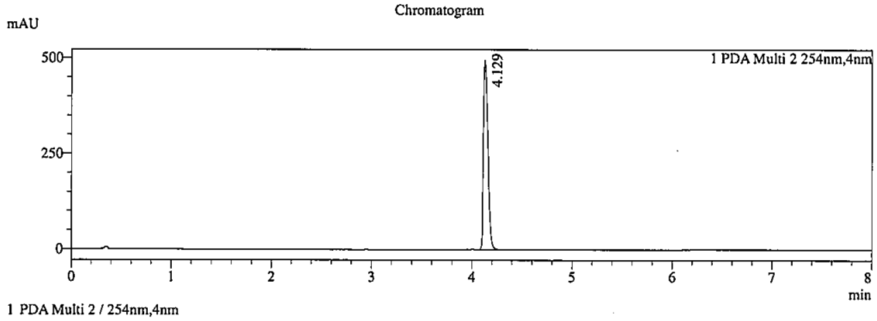

Integration Result

Peak Table

PDA Ch2 254nm

| Peak# | Ret. Time | Height | Height% | USP Width | Area | Area% |
|---|---|---|---|---|---|---|
| 1 | 4.129 | 484289 | 100.000 | 0.084 | 1542826 | 100.000 |

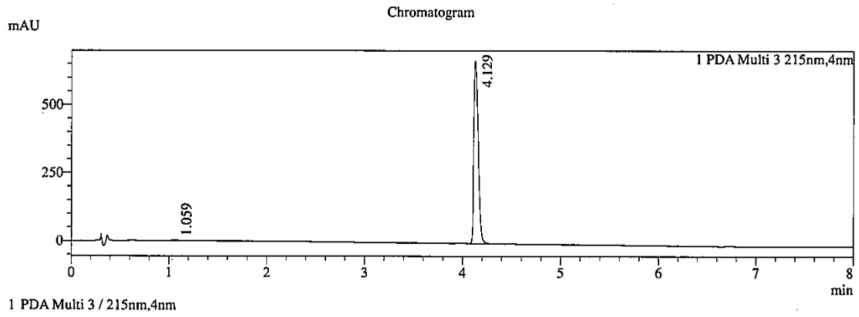

Integration Result

Peak Table

PDA Ch3 215nm

| Peak# | Ret. Time | Height | Height% | USP Width | Area | Area% |
|---|---|---|---|---|---|---|
| 1 | 1.059 | 3033 | 0.461 | 0.071 | 8450 | 0.401 |
| 2 | 4.129 | 655545 | 99.539 | 0.084 | 2099153 | 99.599 |

HPLC chromatogram of compound **3**

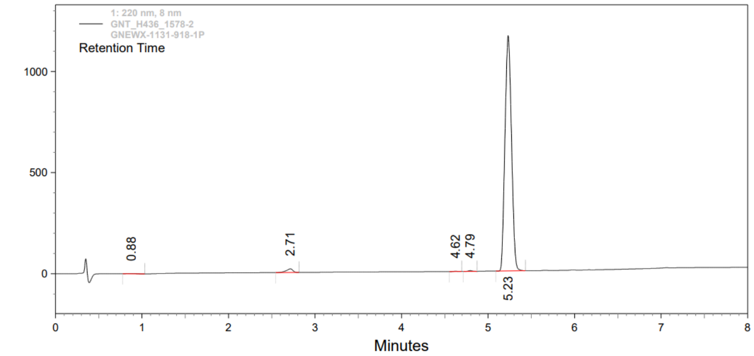

***1: 220 nm, 8 nm***

| *Retention Time* | *Height* | *Area* | *Area Percent* |
|---|---|---|---|
| 0.88 | 2006 | 14264 | 0.23 |
| 2.71 | 17825 | 94685 | 1.52 |
| 4.62 | 1574 | 5119 | 0.08 |
| 4.79 | 4288 | 13870 | 0.22 |
| 5.23 | 1156786 | 6087427 | 97.94 |

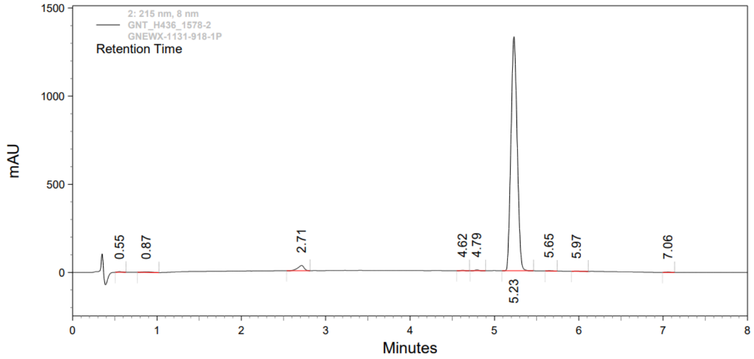

***2: 215 nm, 8 nm***

| *Retention Time* | *Height* | *Area* | *Area Percent* |
|---|---|---|---|
| 0.55 | 3225 | 12600 | 0.18 |
| 0.87 | 3232 | 23433 | 0.33 |
| 2.71 | 29397 | 156299 | 2.23 |
| 4.62 | 2484 | 7826 | 0.11 |
| 4.79 | 4768 | 15348 | 0.22 |
| 5.23 | 1320766 | 6789788 | 96.69 |
| 5.65 | 1640 | 5918 | 0.08 |
| 5.97 | 1527 | 5675 | 0.08 |
| 7.06 | 1670 | 5605 | 0.08 |

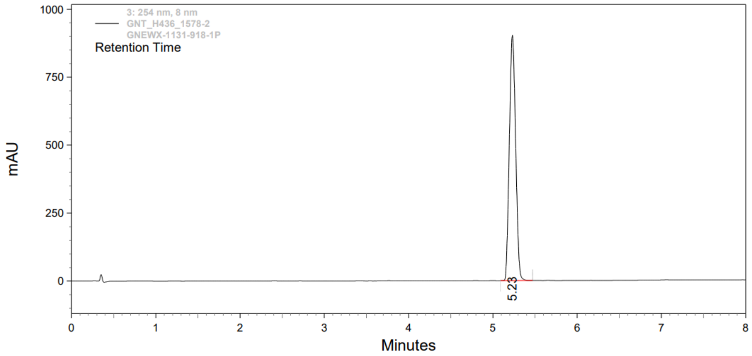

***3: 254 nm, 8 nm***

| *Retention Time* | *Height* | *Area* | *Area Percent* |
|---|---|---|---|
| 5.23 | 897527 | 4692528 | 100.00 |

HPLC chromatogram of compound **4**

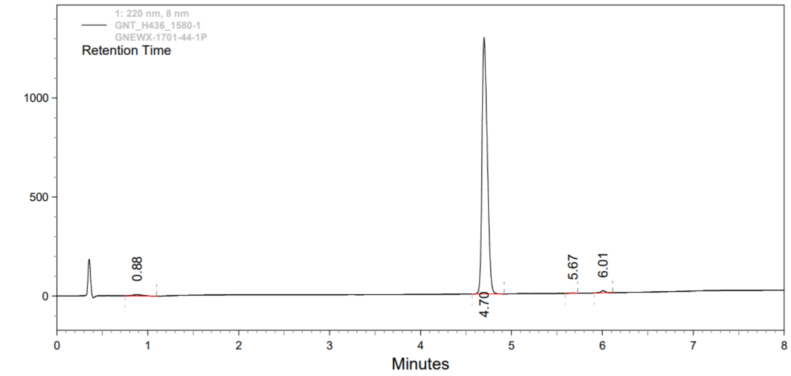

***1: 220 nm, 8 nm***

| *Retention Time* | *Height* | *Area* | *Area Percent* |
|---|---|---|---|
| 0.88 | 5778 | 48393 | 0.91 |
| 4.70 | 1288689 | 5232399 | 98.25 |
| 5.67 | 1644 | 5558 | 0.10 |
| 6.01 | 10759 | 39356 | 0.74 |

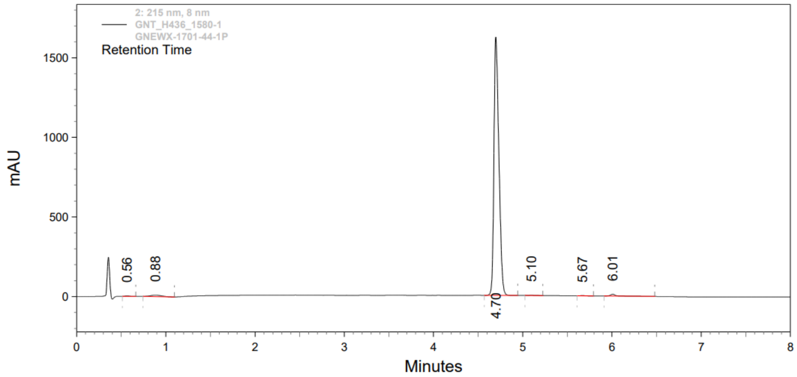

***2: 215 nm, 8 nm***

| *Retention Time* | *Height* | *Area* | *Area Percent* |
|---|---|---|---|
| 0.56 | 2946 | 9287 | 0.14 |
| 0.88 | 9264 | 78075 | 1.22 |
| 4.70 | 1604066 | 6253624 | 97.59 |
| 5.10 | 1747 | 8500 | 0.13 |
| 5.67 | 1685 | 5554 | 0.09 |
| 6.01 | 11358 | 52842 | 0.82 |

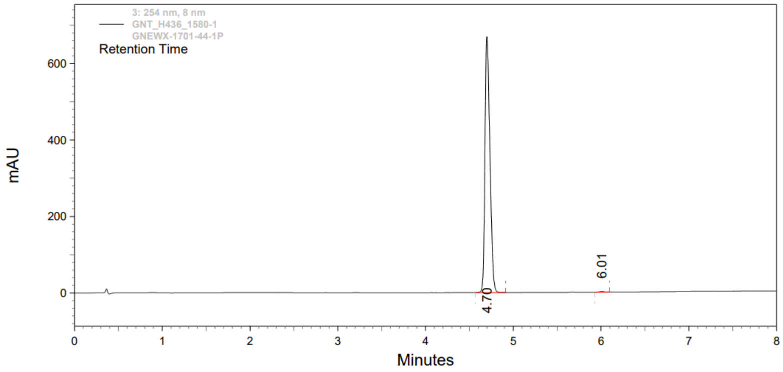

***3: 254 nm, 8 nm***

| *Retention Time* | *Height* | *Area* | *Area Percent* |
|---|---|---|---|
| 4.70 | 665091 | 2626651 | 99.73 |
| 6.01 | 1968 | 7007 | 0.27 |

HPLC chromatogram of compound **5**

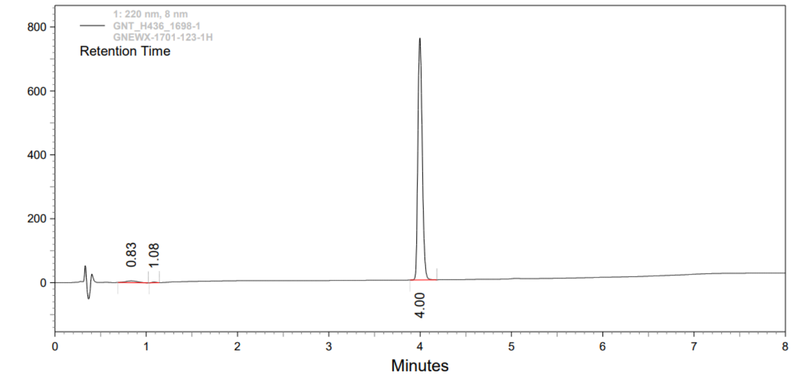

***1: 220 nm, 8 nm***

| *Retention Time* | *Height* | *Area* | *Area Percent* |
|---|---|---|---|
| 0.83 | 5245 | 47427 | 1.85 |
| 1.08 | 2110 | 6804 | 0.27 |
| 4.00 | 753425 | 2510118 | 97.89 |

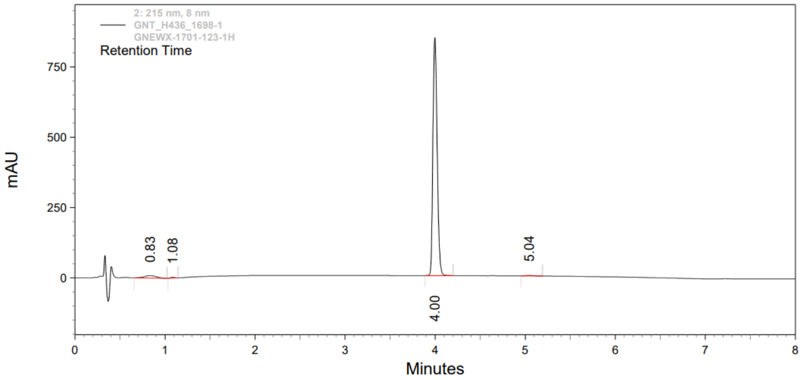

***2: 215 nm, 8 nm***

| *Retention Time* | *Height* | *Area* | *Area Percent* |
|---|---|---|---|
| 0.83 | 8794 | 83696 | 2.90 |
| 1.08 | 3010 | 9655 | 0.33 |
| 4.00 | 842279 | 2784306 | 96.38 |
| 5.04 | 1857 | 11275 | 0.39 |

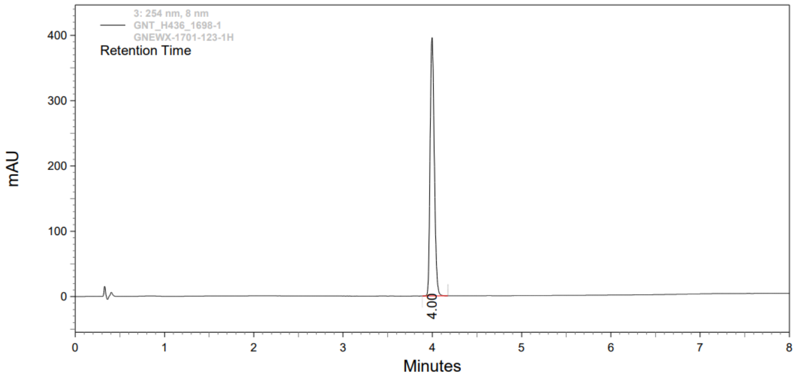

***3: 254 nm, 8 nm***

| *Retention Time* | *Height* | *Area* | *Area Percent* |
|---|---|---|---|
| 4.00 | 393524 | 1296194 | 100.00 |
