## Supplementary figures and images for "CryoEM reveals unprecedented binding site for Na_V_1.7 inhibitors enabling rational design of potent hybrid inhibitors"

### Supplementary Fig 1

#### Supplementary Figure 1

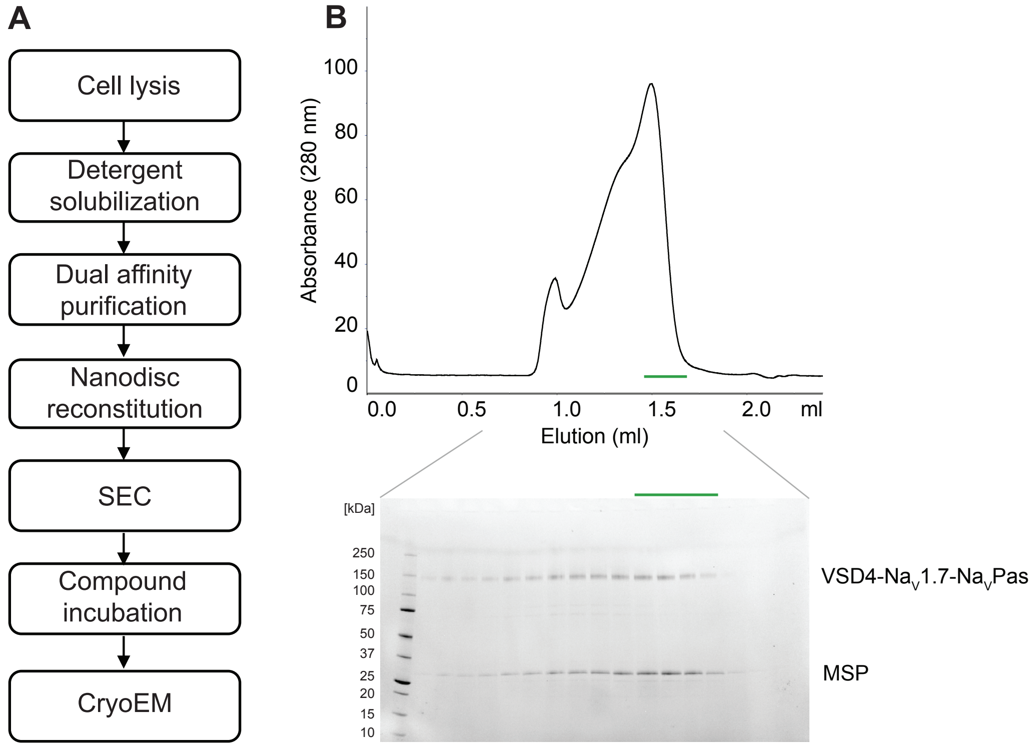

### Supplementary Fig 2

#### Supplementary Figure 2

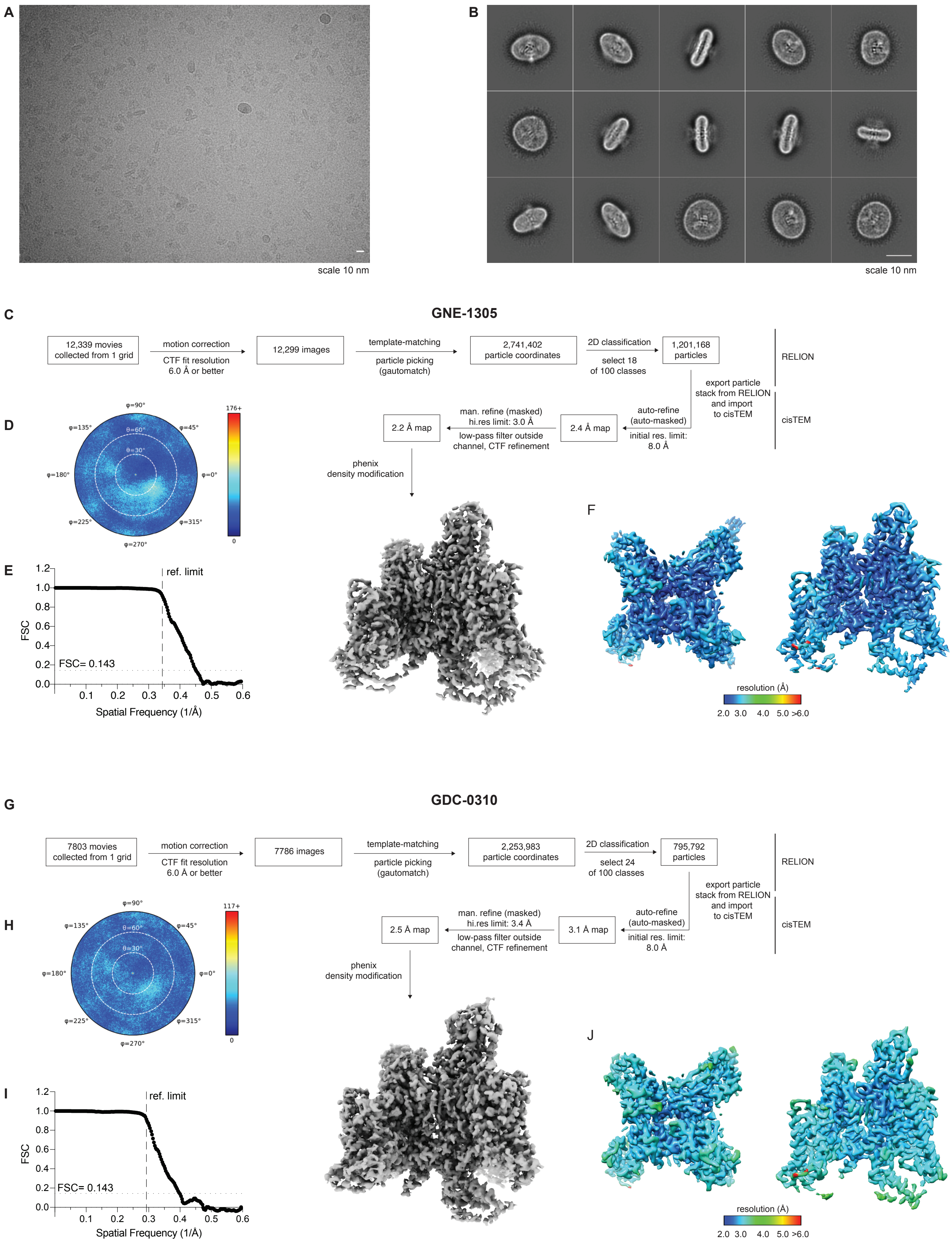

### Supplementary Fig 3

Supplementary Figure 3

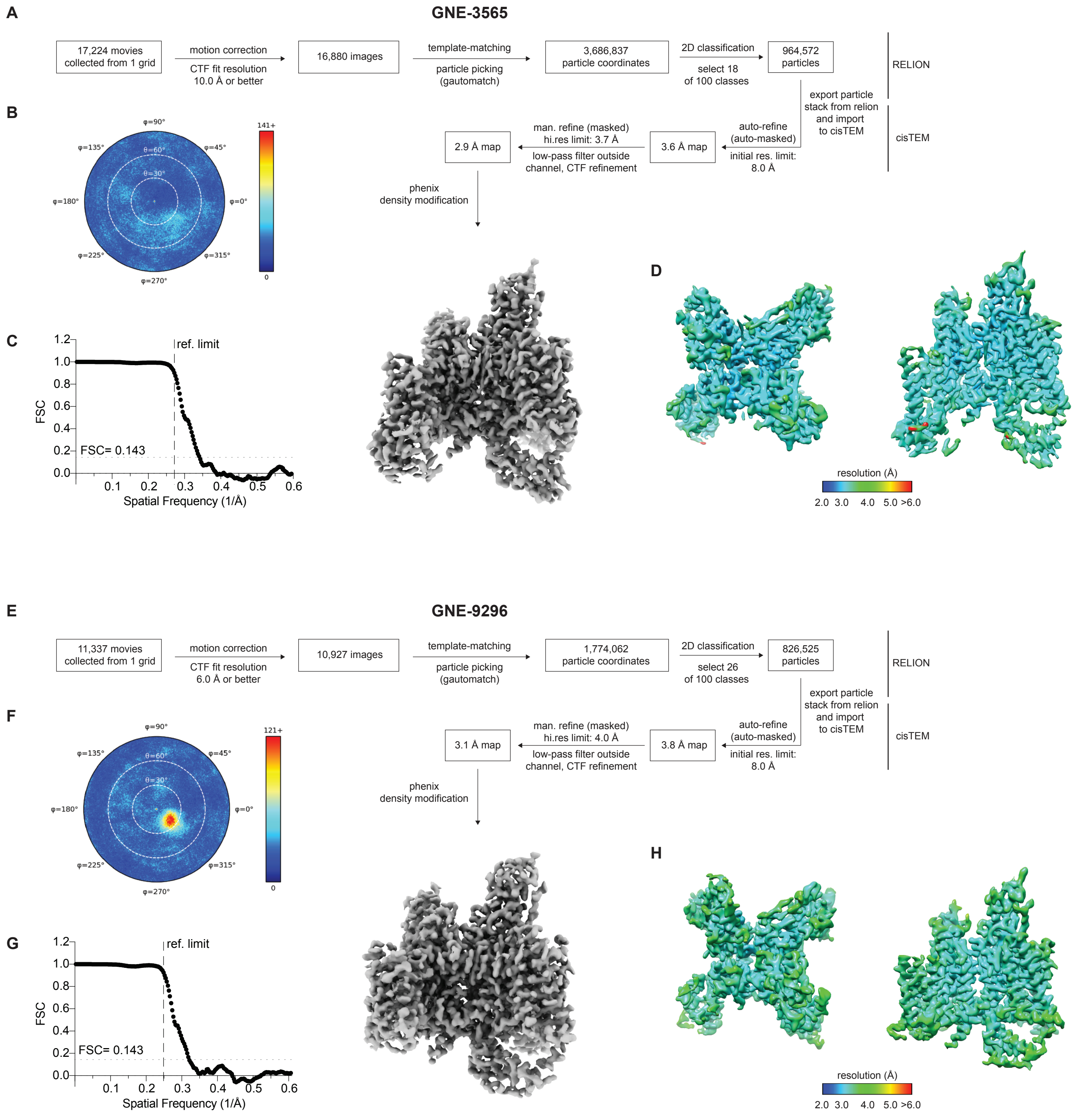

### Supplementary Fig 4

Supplementary Figure 4

A

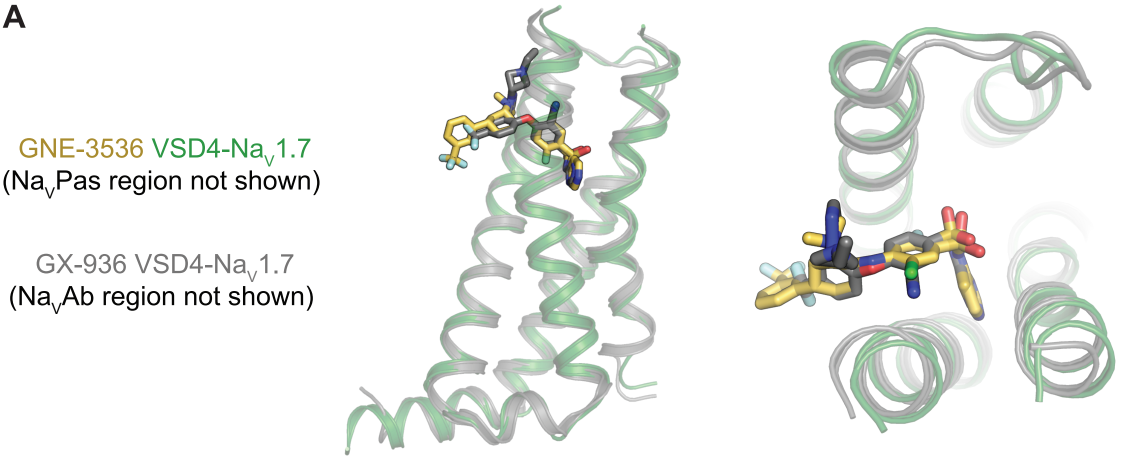


B

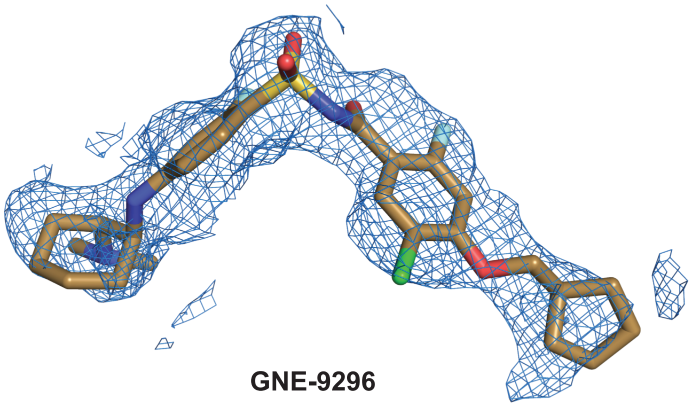


C

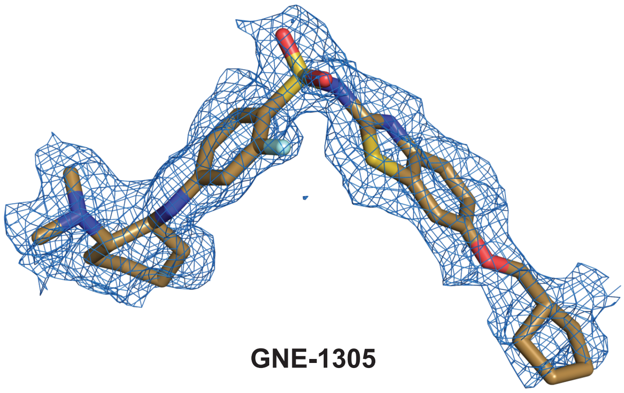
